# Cathepsin B regulates ovarian reserve quality and quantity via mitophagy by modulating IGF1R turnover

**DOI:** 10.1101/2024.02.14.580410

**Authors:** Aradhana Mohanty, Anjali Kumari, Lava Kumar. S, Ajith Kumar, Pravin Birajdar, Rohit Beniwal, Mohd Athar, Kiran Kumar P, H.B.D. Prasada Rao

## Abstract

The quality and quantity of the ovarian reserve are meticulously regulated through various cell death pathways to guarantee the availability of high-quality oocytes for fertilization. While apoptosis is recognized for contributing to maintaining ovarian reserve, the involvement of other cell death pathways remains unclear. Employing chemical genetics and proteomics, this study reveals the crucial involvement of Cathepsin B in maintaining the ovarian reserve. Results indicate that apoptosis and autophagy play pivotal roles, and inhibiting these pathways significantly increases follicle numbers. Proteomics reveals a dynamic shift from apoptosis to autophagy during follicular development, with Cathepsin B emerging as a key player in this transition. Inhibiting Cathepsin B not only mimics the augmented oocyte reserve observed with autophagy inhibition but also upregulated IGF1R and AKT-mTOR pathways without compromising fertility. Further, IGF1R inhibition partially compromised the protective effects of Cathepsin B inhibition on oocyte reserves, suggesting their interdependence. This association is further supported by the finding that Cathepsin B can degrade IGF1R in vitro. Moreover, the increased IGF1R levels enhance the oocyte mitochondrial membrane potential via transcriptional regulation of mitochondrial biogenesis and mitophagy genes. Remarkably, this Cathepsin B-dependent ovarian reserve maintenance mechanism is conserved in higher-order vertebrates. Cumulatively, our study sheds valuable light on the intricate interplay of autophagy, Cathepsin B, and growth factors in ovarian reserve maintenance, offering potential implications for fertility research.

## Introduction

Throughout a female’s reproductive lifespan, regulated cell death mechanisms play fundamental roles at various stages of ovarian development, like follicle formation, development, ovulation, and eliminating damaged oocytes ^1–8^. For example, in humans, the number of oocytes follows a continuous decreasing trend from a peak of 6-7 million oocytes at 20 weeks of gestation to less than 1 million at birth. As a girl reaches menarche, the number further decreases to approximately 300,000-400,000 oocytes ^7, 9–13^. During the reproductive years, the rate of decline remains relatively steady at about 1,000 follicles per month, but this decline accelerates significantly after the age of approximately 37 years. Ultimately, at menopause, the number of remaining follicles drops below 1,000, indicating that ovarian reserve dictates the reproductive lifespan in females ^9, 10, 12^. This pattern of oocyte attrition is not exclusive to humans and is conserved across different species ^8^. While the loss of germ cells is ongoing throughout a female’s life, significant loss occurs with early oocytes called primordial and primary follicles^4, 10^. It is widely acknowledged that oocyte apoptotic cell death processes cause atresia in primordial and primary follicles via the DNA damage response pathway (DDR) ^8, 14^. DDR in early oocytes before birth resembles that observed in somatic cells. It involves the initiation of checkpoint signaling through activating ATM/ATR kinases, which in turn phosphorylate downstream targets to enhance DDR signaling, support damage repair, or trigger programmed cell death. Following birth, primordial follicles constitutively express an alternative member of the p53 family, TAp63, making them highly susceptible to damage-induced apoptosis ^15–19^.

On the other hand, active autophagy was observed in perinatal ovaries consisting of very early oocytes ^20^. Inhibition of autophagy increased the number of cyst oocytes and delayed the folliculogenesis in organ cultures^20^. In addition, depletion of Becn1 or Atg7 resulted in elevated germ cell loss and subfertility, indicating the roles of other cell death pathways in oocyte reserve maintenance ^21^. However, the significance of alternative death pathways and their mechanisms in early follicular atresia is unknown. This study shows that inhibition of lysosomal cysteine protease Cathepsin B enhances the oocyte reserve via regulating IGF1R turnover.

## Results

### Postnatally, both apoptosis and autophagy play integral roles in preserving ovarian reserve

In exploring the fundamental pathways contributing to early ovarian follicle depletion, we thoroughly investigated specific inhibitors administered to 5-day-old female mice. This particular stage was chosen due to the abundance of early follicles, allowing for the administration of inhibitors without causing harm to the mice. The experimental cohorts were subjected to a dosage of 5mg/kg body weight for an apoptosis inhibitor (Z-VAD-FMK) 50mg/kg for an autophagy inhibitor (3-MA), 10mg/kg for a pyroptosis inhibitor (Disulfiram), and 5mg/kg for a necrosis inhibitor (GSK^’^ 872), with a single dose of alternate day administration (Fig.1a)^22–25^. Ovaries were collected on the 10th day, subsequently fixed, sectioned, and subjected to immunostaining for the germ cell-specific marker, mouse vasa homologue (MVH), and the guardian of germ cells, TAp63α (Fig. 1b). Analysis of follicle counts unveiled that mice treated with apoptosis, autophagy, and pyroptosis inhibitors exhibited 2.4, 2.4, and 1.7-fold increases in ovarian follicle numbers, respectively, compared to the vehicle control group (Fig.1c and Fig.S1 a, b and c). In contrast, mice treated with the necrosis inhibitor displayed no significant variations, suggesting that apoptosis and autophagy hold equal importance in the early loss of ovarian follicles, while pyroptosis may have a secondary impact (Fig.1c and Fig.S1 a, b and c). Notably, the co-administration of inhibitors targeting apoptosis and autophagy from 5 to 10 days postpartum (dpp), or exclusively using inhibitors for apoptosis, autophagy, pyroptosis, and necrosis within a 5 to 21-day timeframe, resulted in lethality in mice, underscoring the intricate balance these pathways uphold for the overall health of the animals.

**Fig. 1:**
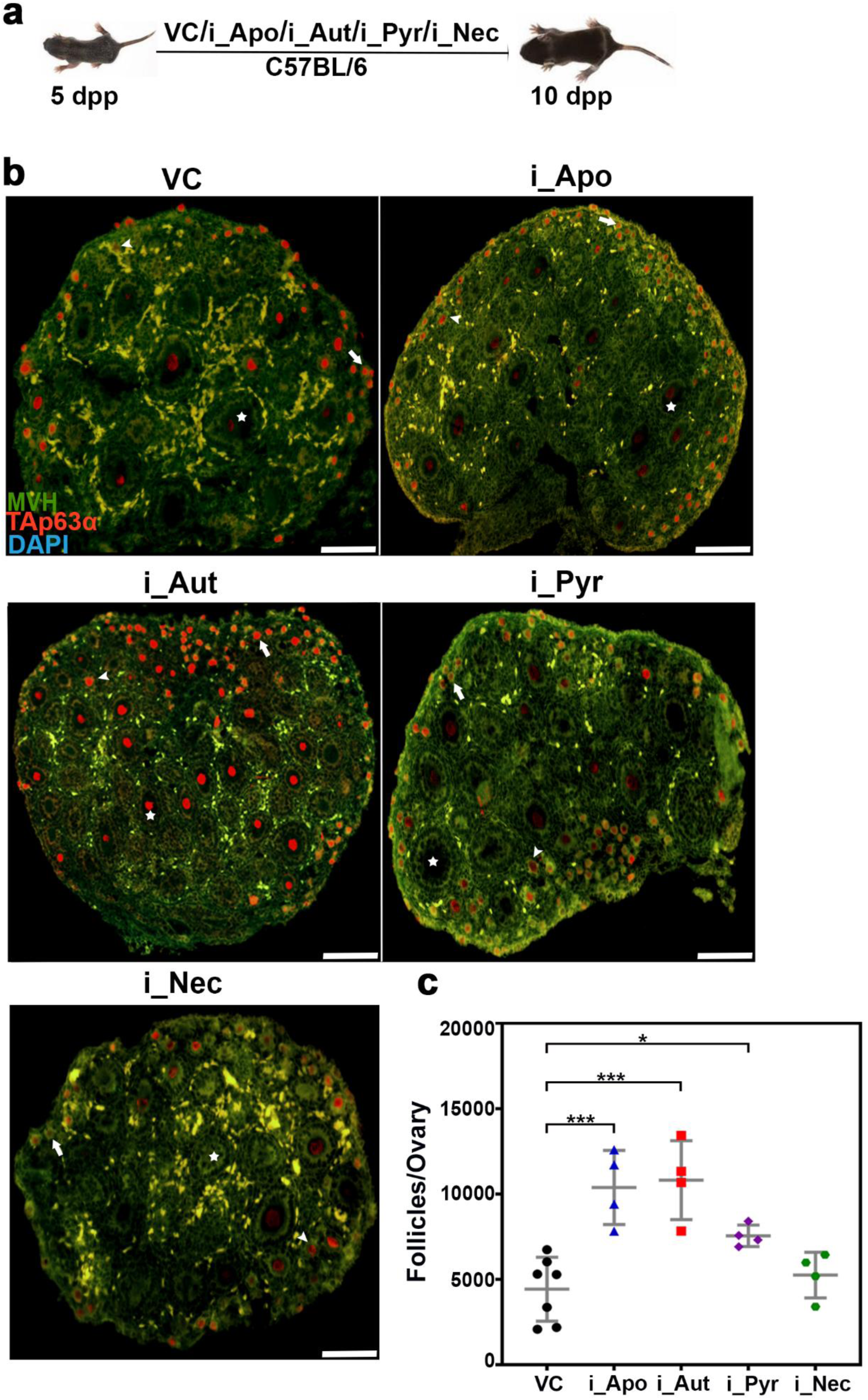
Role of death pathways in ovarian reserve maintenance. **a,** Intraperitoneal injection of death pathway inhibitors: an experimental regimen. **b,** 10 days postpartum (dpp) ovary sections from VC, i_Apo, i_Aut, i_Pyr, and i_Nec -treated females immunostained for TAp63α (Red), MVH (green) and DNA (blue). **c,** Follicle counts at 10 dpp from VC, i_Apo, i_Aut, i_Pyr, and i_Nec - treated females. VC, vehicle control; i_Apo, an inhibitor of apoptosis; i_Aut, an inhibitor of autophagy; i_Pyr, an inhibitor of pyroptosis; i_Nec, an inhibitor of necrosis. Arrows, arrowheads, and stars highlight the primordial, primary, and secondary follicles, respectively. \*\*\**P* ≤ 0.001, \**P* ≤ 0.01, *ns ≥* 0.1 unpaired t test. Error bars show mean ± SD. Non-significance P values are not represented in the figure. Scale bars for ovary sections are 100 μm.

### Autophagy and lysosomal cysteine proteases are upregulated during folliculogenesis

To comprehend the intricacies of death pathways in folliculogenesis, we conducted untargeted proteomics on ovaries from mice at different developmental stages (5, 10, and 21 days postpartum). The results revealed the identification of 4914, 4432, and 4843 proteins in 5, 10, and 21-day-old ovaries (n=3). Intriguingly, 70, 5, and 48 proteins were uniquely identified in 5, 10, and 21-day-old ovaries, respectively, while 4529 proteins were common across all samples (Fig.2a). Multivariate discriminative analysis unveiled a significant separation between the three developmental stages, with 25.3% PC1 and 18.1% PC2 contributing to the distinction (Fig.2b). The generated heat map and hierarchical clustering analysis demonstrated distinct clusters corresponding to 5, 10, and 21-day-old ovaries (Fig.2c). Delving into the biological functions of the identified proteins, we observed a downregulation of apoptosis-related genes in 10-day-old ovaries compared to 5-day-old ovaries. Concurrently, upregulation of autophagy-related genes in 10-day-old ovaries suggested the activation of autophagy during this developmental phase, coinciding with decreased apoptosis activity (Fig.2d). A similar comparison between 21 and 10-day-old ovaries indicated an upregulation of autophagy-related genes, signifying heightened autophagic activity in 21dpp (Fig.2e). The balanced regulation of apoptosis genes in 21-day-old ovaries, compared to 10-day-old ovaries, suggested intricate dynamics, possibly influenced by other somatic cells like cumulus cells (Fig.2d and e). Corroborating the proteomic findings, Western blot analyses validated reduced DNA damage levels (gH2AX), diminished apoptosis levels (cc3), decreased oocyte reserve (TAp63α), and increased autophagy levels (LC3 and P62) in ovaries at 5, 10, and 21 days postpartum (Fig.2f). Notably, the elevated autophagy levels during folliculogenesis prompted an in-depth exploration of the abundance of proteins at 5 dpp compared to 10 dpp and 10 dpp compared to 21 dpp. Among the top 50 most abundant proteins, significant findings included the presence of cysteine proteases Cathepsin B and D and the autophagy protein ATG7 in ovaries at both 10 and 21 days of age (Fig.2g and h). Subsequent validation through Western blots conclusively confirmed a discernible upregulation of lysosomal cysteine proteases and autophagy during folliculogenesis (Fig.2i).

**Fig. 2:**
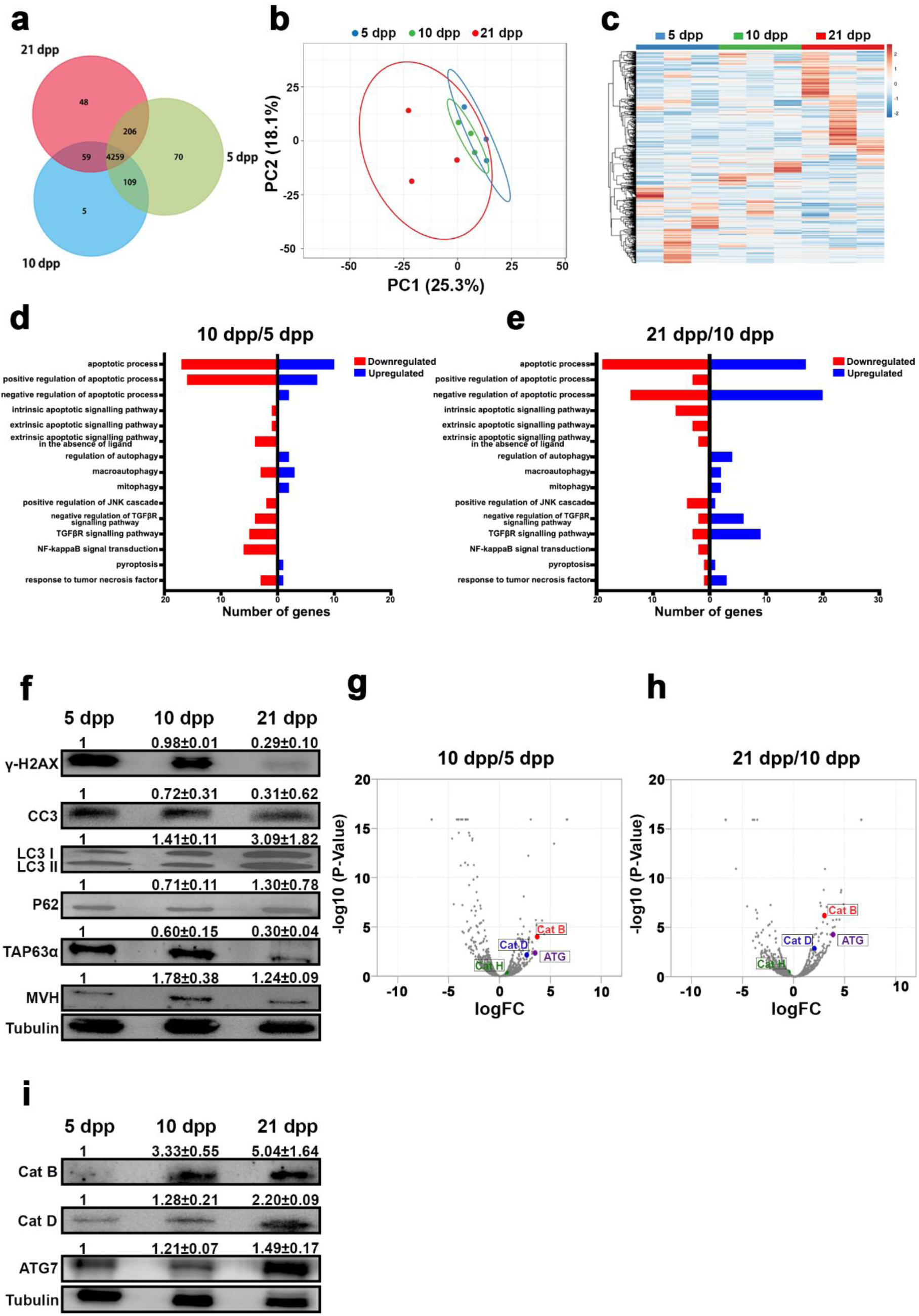
Proteomic profiling of mice ovary in different age groups. **a,** Venn diagram of 5,10, and 21 dpp mice ovaries proteome. **b**, Principal Component Analysis (PCA) shows the difference between the 5,10, and 21 dpp proteome profiles. **c,** The relative abundance of differential proteins in 5,10, and 21 dpp is presented as a heatmap. **d and e,** Distribution of death pathways related proteins identified in 5 and 10 dpp ovaries, 10 and 21 dpp ovaries, respectively. **f,** Western blot analysis of γH2AX, cc3, LC3, P62, TAp63α, MVH and Tubulin from 5, 10, and 21 dpp ovaries. **g and h,** A volcano plot showing *P* values versus fold changes of all proteins in 5 and 10 dpp ovaries, 10 and 21 dpp ovaries, respectively. **i,** Western blot analysis of Cat B, Cat D, ATG7, and Tubulin from 5, 10, and 21 dpp ovaries.

### Cathepsin B inhibition protects the ovarian reserve

Since Cathepsin B and D are abundant in the 10 and 21 dpp ovaries, previous results in ***C.*** *elegance* show that the Insulin signaling pathway regulates the quality of oocytes via Cathepsin B activity ^26^. Therefore, to know the effect of Cathepsin B and D on ovarian reserves, we pharmacologically inhibited Cathepsin B and D. Administration of Cathepsin B inhibitor Myricetin 37.5mg/kg body weight and Cathepsin D inhibitor Pepstatin A 20mg/kg body weight to 10-day-old female mice daily with a single dose and collected the ovaries on the 21st day (Fig. 3a)^27, 28^. Ovaries were fixed, sectioned, and immunostained for TAp63α (Fig.3b). Follicle counts revealed that Cathepsin B-inhibited ovaries contained 2.4 times more follicles than control ovaries. At the same time, Cathepsin D-inhibited ovaries do not exhibit an appreciable rise in follicles compared to the control (Fig.3c). Moreover, the follicular analysis showed an increase in early follicles, such as primordial, primary and secondary follicles, in Cathepsin B-inhibited ovaries (Fig.3d). Furthermore, the follicle protective effect by Cathepsin B inhibition was dose dependent (Fig.S2b). Subsequent western blot analysis confirmed that the inhibition of Cathepsin B resulted in 45 and 33% reduction in the protein levels of Cathepsin B and LC3 respectively. In contrast, TAp63α levels increased 4-fold, indicating that Cathepsin B inhibition protects ovarian reserve (Fig3e). To assess whether the protected oocytes can undergo maturation and embryogenesis, mice were treated with the Cathepsin B inhibitor from 10 to 21 dpp and sacrificed on the 60th day (Fig.S2d). Serum and oocytes were collected for endocrine profiling and assessment of oocyte quality (Fig.S2d). Results indicated no significant differences in oocyte maturation and embryogenesis, suggesting that the protected oocytes are of high quality (Fig.S2f-i). Endocrine profiling revealed no discernible impact of Cathepsin B treatments (Fig.S2e). Additionally, a cohort of treated animals were monitored for up to 8 litters, and no differences in litter size or fertility were observed compared to controls, indicating that Cathepsin B inhibitor treatment does not affect fertility (Fig.S2j).

**Fig. 3:**
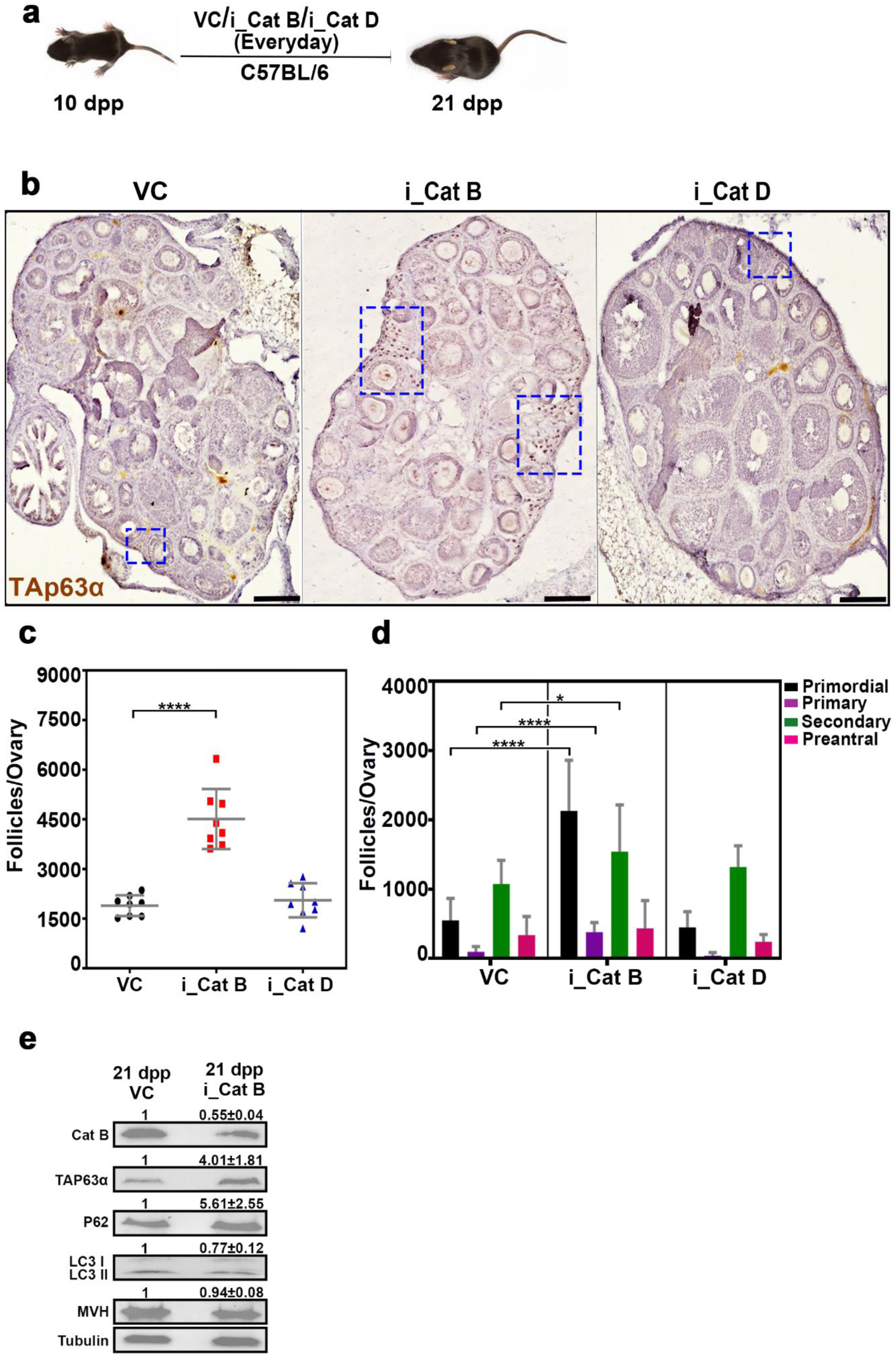
Inhibition of Cathepsin B protects the ovarian reserve. **a,** Experimental regime for intraperitoneal injection of Cathepsin B and D inhibitors. **b,** 21 dpp ovary sections from VC, i_Cat B, and i_Cat D -treated females immunostained for TAp63α with hematoxylin counterstaining. **c,** Follicle counts at 21 dpp from VC, i_Cat B, and i_Cat D -treated females. **d,** Types of follicles from 21 dpp from VC, i_Cat B, and i_Cat D -treated females. **e,** Western blot analysis of Cat B, TAp63α, P62, LC3, MVH, and Tubulin from VC, and i_Cat B -treated ovaries. VC, vehicle control; i_Cat B, an inhibitor of Cathepsin B; i_Cat D, an inhibitor of Cathepsin D. The blue square indicates the follicle-enriched areas. \*\*\*\**P* ≤ 0.0001, \**P* ≤ 0.05, *ns ≥* 0.1 unpaired t test. Error bars show mean ± SD. Non-significance P values are not represented in the figure. Scale bars for ovary sections are 100 μm.

To evaluate the sustained protective effects of Cathepsin B inhibition on ovarian reserve, we employed a daily administration of the Cathepsin B inhibitor from postnatal day 10 to 21, followed by a 6-month observation period (Fig.S2a). Subsequently, ovaries were collected, fixed, sectioned, and subjected to immunostaining for TAp63α. Examination of follicular quantification revealed a 1.9-fold increase in the ovaries treated with the Cathepsin B inhibitor compared to the control group (Fig.S2c). This increase was predominantly observed in enhanced primordial and primary follicles, suggesting that early Cathepsin B inhibition may contribute to the long-term protection of the follicular reserve.

Furthermore, in assessing the genetic prerequisites for early follicle protection through Cathepsin B inhibition, we conducted co-administration experiments with inhibitors targeting apoptosis, autophagy, pyroptosis, or necrosis from 5 to 10 days dpp (FigS3a). Ovaries were collected on the 10th day, fixed, sectioned, and subjected to immunostaining for MVH and TAp63α. Our analysis revealed a reduction in follicle numbers when the Cathepsin B inhibitor was co-administered with the apoptosis inhibitor, in contrast to the use of Cathepsin B or apoptosis inhibitor alone, suggesting an independent functioning of the two pathways and lethality to follicles when both death pathways are inhibited. Similarly, co-administration of the Cathepsin B inhibitor with the autophagy inhibitor replicated the effects of autophagy inhibitor treatment, indicating the involvement of Cathepsin B in regulating the ovarian reserve through the autophagy pathway. No significant difference in follicle numbers was observed when the Cathepsin B inhibitor was co-administered with the pyroptosis inhibitor, suggesting redundant roles in this context. Finally, co-administration of the Cathepsin B inhibitor with the necrosis inhibitor resulted in a decrease in follicle numbers compared to Cathepsin B alone, indicating a potential role of necrosis in Cathepsin B-inhibited follicles (Fig.S3b).

### Cathepsin B regulates IGF-1R turnover

Besides, we conducted untargeted proteomics on ovaries from 21-day-old control and Cathepsin B inhibitor-treated mice ovaries to elucidate the protective mechanism of Cathepsin B inhibition on the ovarian reserve. Our analysis revealed the identification of 4274 and 4183 proteins in control and Cathepsin B inhibited ovaries, respectively (n=3). Notably, 127 and 36 proteins were found exclusively in control and Cathepsin B-inhibited ovaries, while 4147 proteins were shared between both groups (Fig.4a). Multivariate discriminative analysis highlighted a significant separation between the two samples, with 31.1% PC1 and 24.6% PC2 contributing to this distinction (Fig.4b). The subsequent heat map and hierarchical clustering analysis confirmed distinct clusters corresponding to control and Cathepsin B inhibited ovaries (Fig.4c). Examining the biological functions of the identified proteins, we observed a downregulation of autophagy-related genes and an upregulation of insulin-regulatory genes in Cathepsin B inhibited ovaries compared to the control (Fig.4d). The elevated insulin regulatory pathways in Cathepsin B inhibited ovaries, prompted a detailed investigation into the protein abundance. Among the top 50 most abundant proteins, a significant finding was the presence of IGF1R (Fig.4e). Substantiating the proteomic results, the Western blot analyses confirmed increased levels of IGF1, IGF1R, and AKT pathway proteins indicating that Cathepsin B plays a role in regulating the insulin signaling pathway (Fig.4g).

**Fig. 4:**
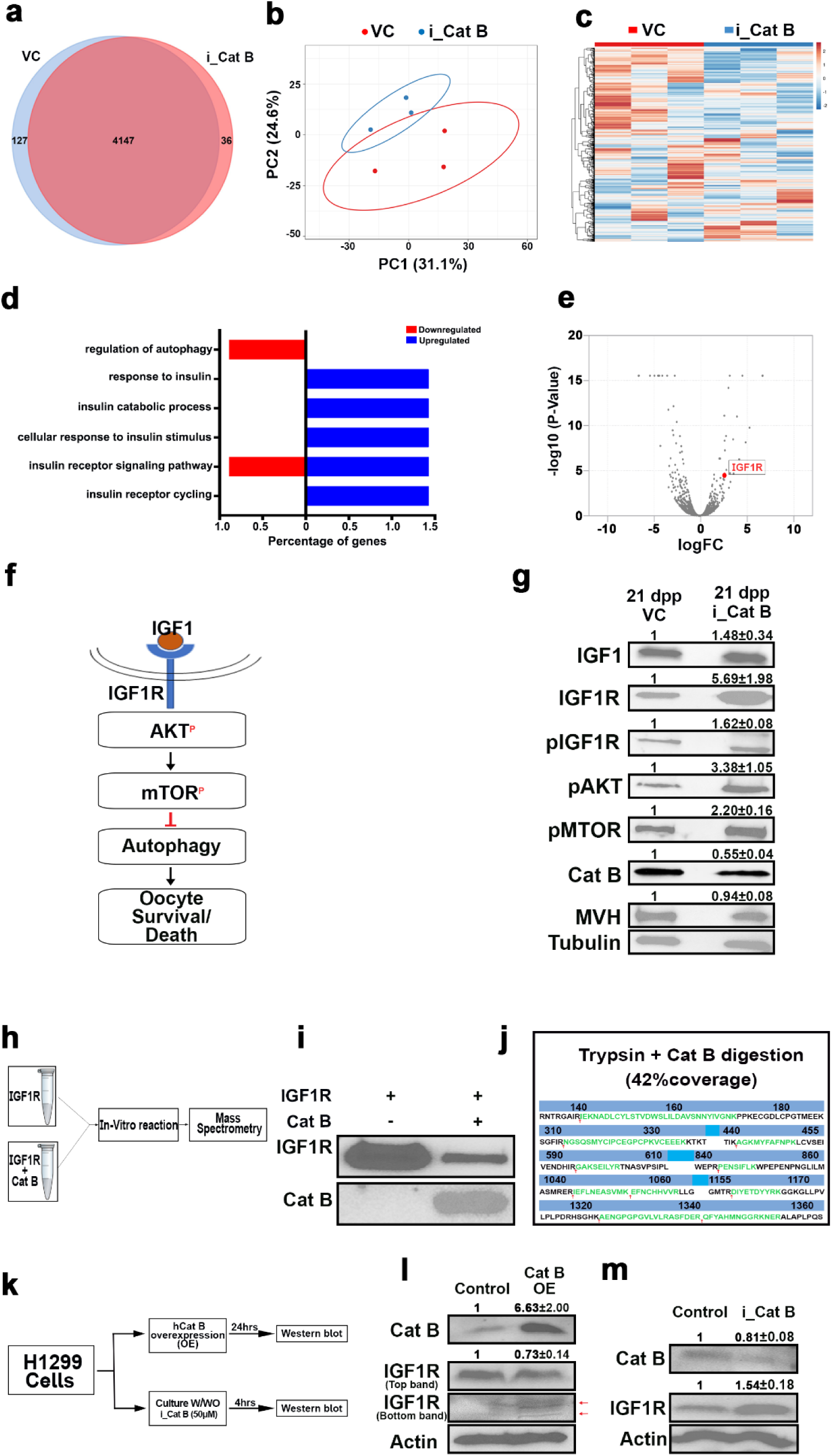
Proteomic profiling of Control and i_Cat B treated mice ovaries. **a,** Venn diagram of VC and i_Cat B -treated ovaries proteome. **b**, Principal Component Analysis (PCA) shows the difference between the VC and i_Cat B -treated ovaries proteome profiles. **c,** Relative abundance of differential proteins in VC and i_Cat B -treated ovaries presented as a heatmap. **d,** Distribution of all protein classes identified in VC and i_Cat B - treated ovaries according to biological process. **e,** A volcano plot showing *P* values versus fold changes of all proteins in VC and i_Cat B -treated ovaries. **f**, Schematic representation of insulin signalling pathway in oocytes. **g,** Western blot analysis of IGF1, IGF1R, pIGF1R, pAKT, pMTOR, Cat B, MVH and Tubulin from VC and i_Cat B -treated ovaries. **h**, Experimental regimen for in vitro reaction followed by mass spectrometry. **i**, Western blot analysis of in vitro digestion by Cat B. **j**, Schematic representation of invitro digested peptides of IGF1R identified in mass spectrometry. **k**, Experimental regimen for H1299 cell culture experiments. **l,** Western blot analysis of Cat B, IGF1R, and Actin from Control and Cat B-overexpressed H1299 cells. **m,** Western blot analysis of Cat B, IGF1R, and Actin from Control and i_Cat B -treated H1299 cells. VC, vehicle control; i_Cat B, an inhibitor of Cathepsin B.

Specifically, we observed a 5.7-fold accumulation of IGF1R levels in ovaries inhibited with Cathepsin B, leading us to consider the potential of IGF1R being a substrate for the cysteine protease Cathepsin B (Fig.4g). To explore this hypothesis, we conducted an in vitro reaction involving human recombinant Cathepsin B incubated with recombinant IGF1R. Subsequently, we quenched the reaction and subjected it to western blot and mass spectrometry analysis (Fig.4h). The western blot results indicated a reduction in IGF1R levels in the presence of Cathepsin B (Fig.4i). Mass spectrometry analysis further revealed that IGF1R treated with Cathepsin B and trypsin displayed ten specific digestion sites, in contrast to trypsin alone (Fig.4j). To validate whether IGF1R is indeed a substrate for Cathepsin B, we over-expressed human Cathepsin B in H1299 cells (Fig.4k). After 24 hours, analysis of Cathepsin B and IGF1R levels through western blot suggested that the over-expression of Cathepsin B led to a decrease in IGF1R levels, accompanied by observation of specific degraded bands compared to the control (Fig.4l). Conversely, when a Cathepsin B inhibitor was added to H1299 cells and harvested after 4 hours, western blot analysis showed increased IGF1R levels (Fig.4m). These in vitro in vivo results suggest that inhibiting Cathepsin B leads to the accumulation of IGF1R, supporting the notion that IGF1R may act as a substrate for Cathepsin B.

### The inhibition of IGF1R partially mitigates the effect of Cathepsin B inhibition on the ovarian reserve

In investigating the observed preservation of a greater number of follicles in the ovaries of mice with Cathepsin B inhibition and the concurrent elevation of IGF1R levels, we sought to understand by exploring the potential interplay between these factors. To ascertain the comprehensive impact, we conducted a study where we administered a Cathepsin B inhibitor, Myricitin (37.5mg/kg body weight), and an IGF1R inhibitor, PPP (2.5mg/kg body weight), to 10-day-old female mice every alternate day with a single dose (Fig.5a). The objective was to determine whether inhibiting IGF1R in conjunction with Cathepsin B inhibition would diminish ovarian reserve. Ovaries were collected on the 21st day, fixed, sectioned, and immunostained for TAp63α to assess the outcomes of this combined inhibition strategy (Fig.5b). Analysis of follicle counts revealed that Cathepsin B-inhibited ovaries exhibited 1.9 as many follicles as control ovaries. In contrast, ovaries with IGF1R inhibition showed no significant increase in follicles compared to the control group. Intriguingly, mice treated with both Cathepsin B and IGF1R inhibitors displayed a 1.4-fold reduction in follicle count compared to those treated with the Cathepsin B inhibitor alone, suggesting that the increased levels of IGF1R may indeed be responsible for the increased number of follicles in Cathepsin B-inhibited mice ovaries (Fig.5c and d).

**Fig. 5:**
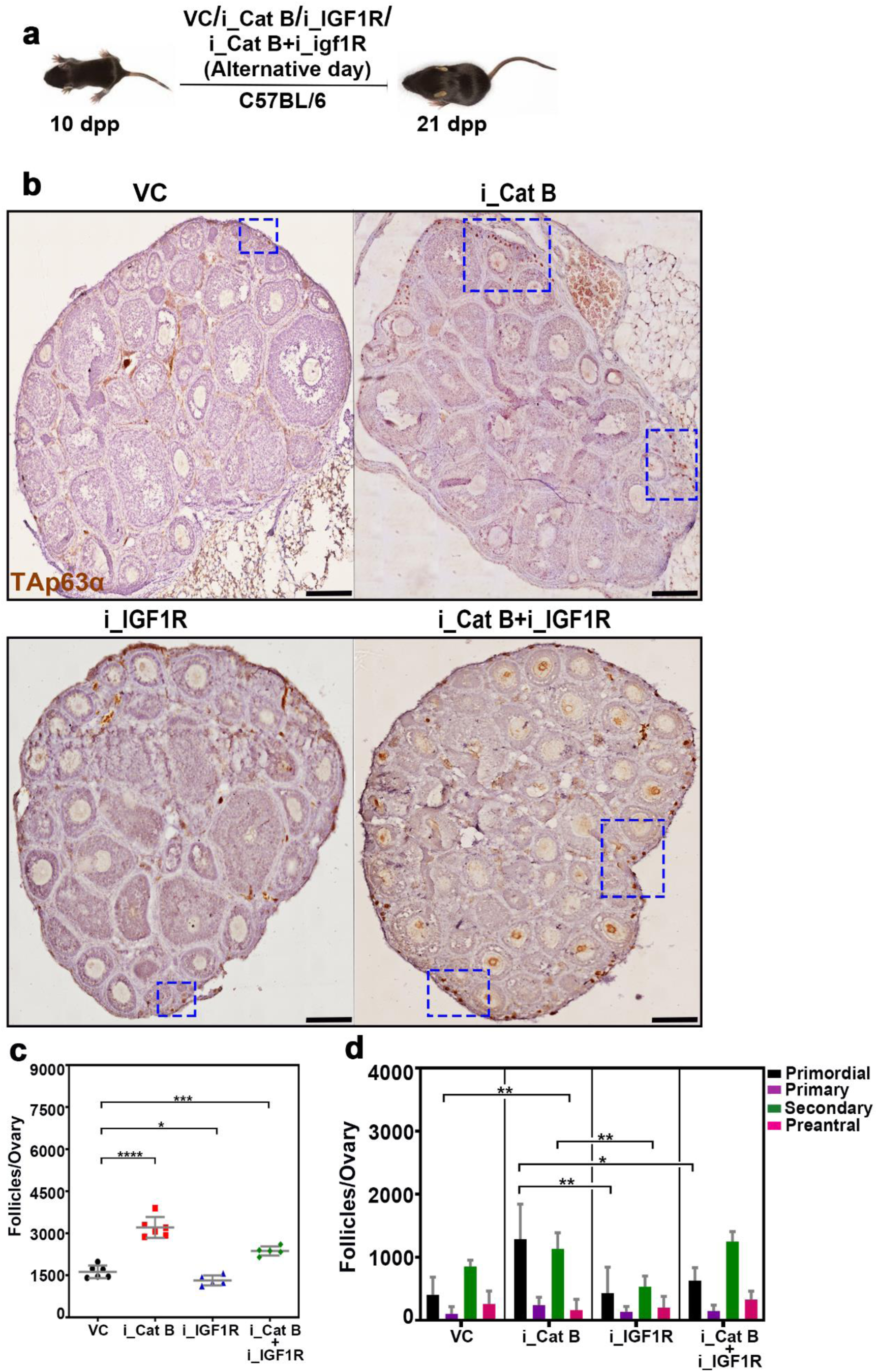
Cathepsin B protects the ovarian reserve via IGF1R. **a,** Experimental regime for intraperitoneal injection of Cathepsin B and IGF1R inhibitors. **b,** 21 dpp ovary sections from VC, i_Cat B, i_IGF1R and i_Cat B + i_IGF1R -treated females immunostained for TAp63α with hematoxylin counterstaining. **c,** Follicle counts at 21 dpp from VC, i_Cat B, i_IGF1R and i_Cat B + i_IGF1R -treated females. **d,** Types of follicles from 21 dpp from VC, i_Cat B, i_IGF1R and i_Cat B + i_IGF1R -treated females. VC, vehicle control; i_Cat B, an inhibitor of Cathepsin B; i_IGF1R, an inhibitor of IGF1R. The blue square indicates the follicle-enriched areas. \*\*\*\**P* ≤ 0.0001, \*\*\**P* ≤ 0.0002, \*\**P* ≤ 0.02, \**P* ≤ 0.04, *ns ≥* 0.1 unpaired t test. Error bars show mean ± SD. Non-significance P values are not represented in the figure. Scale bars for ovary sections are 100 μm.

### Cathepsin B inhibition elevates the mitochondrial membrane potential via enhanced mitophagy

To elucidate the protective mechanism of Cathepsin B inhibition on ovarian reserve through elevated IGF1R levels, we comprehensively analyzed the proteome within IGF1R-associated pathways in both control and Cathepsin B inhibited ovaries. Our findings from the proteome analysis revealed an upregulation of mitophagy and downregulation of mitochondria depolarization in ovaries treated with Cathepsin B, as opposed to the control group (Fig.6a). In addition, previous research on mammalian cells has indicated that IGF1 signaling regulates mitochondrial turnover through mitophagy. Therefore, to delve deeper into understanding how increased mitophagy contributes to the protection of the ovarian reserve, we administered the Cathepsin B inhibitor Myricitin at a dosage of 37.5mg/kg body weight to 10-day-old female mice daily with a single dose. Ovaries were collected on the 21st day, and follicles were isolated and categorized into early and late stages based on their cumulus layers (Fig.6b). Early follicles were identified as those with one layer of cumulus cells, while late follicles had multiple layers. Following the separation, the follicles underwent live staining with JC1 (detecting mitochondrial membrane potential), Mito tracker (for mitochondrial localization), and lysotracker (for lysosomal localization) (Fig. 6c, e and g). Analysis of JC1 in early follicles indicated an increased membrane potential in Cathepsin B treated ovarian follicles compared to controls (Fig.6d). Intriguingly, the mitochondrial localization and intensity were consistent between control and treatment (Fig.6f). However, lysosome activity was elevated in Cathepsin B-treated early follicles compared to the control group, suggesting that elevated mitophagy may clear damaged mitochondria through increased lysosomal activity, contributing to the biogenesis of new mitochondria (Fig.6h). Supporting this notion, we observed elevated levels of reactive oxygen species in control early follicles compared to Cathepsin B inhibited early follicles (Fig.6i-j). We observed similar dynamics in late follicles, however the fold difference is reduced compared to early follicles. Additionally, transcription levels of BNIP3, a key player in mitophagy, and the essential TFAM1 for mitochondrial biogenesis were higher in Cathepsin B treated ovaries than in controls ^29, 30^ (Fig.6k-m). These findings collectively underscore the potential protective role of Cathepsin B inhibition in preserving ovarian reserve through complex regulation of mitophagy and mitochondrial dynamics.

**Fig. 6:**
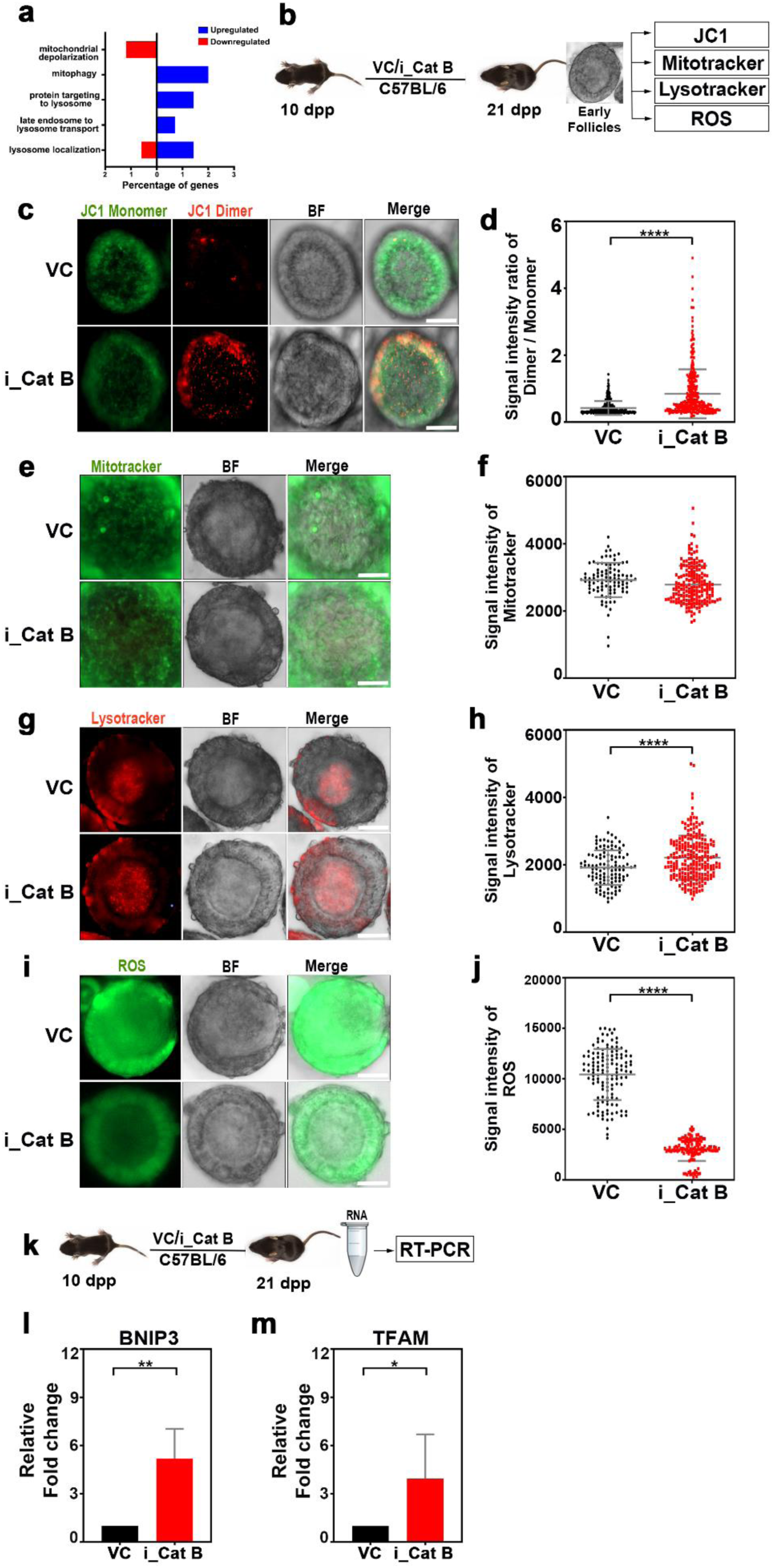
Cathepsin B inhibition regulates mitochondrial dynamics via IGF1R upregulation. **a,** Relative abundance of IGF1R pathway-related proteins in VC and i_Cat B-treated ovaries. **b,** Experimental regime for intraperitoneal injection of VC and i_Cat B -treated ovaries followed by JC1, mitotracker, lysotracker, and ROS experiments. **c,** Early follicles from VC and i_Cat B -treated ovaries were stained for JC1 and imaged at 488nm for JC-1 monomers and 565nm for dimers. **d,** Quantification of the signal intensity ratio of Dimer/Monomer. **e,** Early follicles from VC and i_Cat B -treated ovaries were stained for mitotracker green. **f,** Quantification of the signal intensity of mitotracker. **g,** Early follicles from VC and i_Cat B - treated ovaries were stained for lysotracker red. **h,** Quantification of the signal intensity of lysotracker. **i,** Early follicles from VC and i_Cat B -treated ovaries were stained for ROS. **j,** Quantification of the signal intensity of ROS. **K,** Experimental regime for intraperitoneal injection of VC and i_Cat B -treated ovaries followed by RT PCR experiments. **l and m,** Quantification of the relative fold change of BNIP3, TFAM gene expression in VC and i_Cat B -treated ovaries. VC, vehicle control; i_Cat B, an inhibitor of Cathepsin B. \*\*\*\**P* ≤ 0.0001, \*\**P* ≤ 0.006, \**P* ≤ 0.04, *ns ≥* 0.01 unpaired t test. Error bars show mean ± SD. Non-significance P values are not represented in the figure. Scale bars for follicles are 10 μm.

### Cathepsin B inhibition protects the ovarian reserve in goat ovaries

To investigate the potential protective role of Cathepsin B inhibition on follicular reserve in higher-order vertebrates, we conducted untargeted mass spectrometry analysis on early (1 dpp) and late (10 dpp) goat ovaries. The results revealed the identification of 4327 and 4124 proteins in early and late ovaries (n=2), respectively. Notably, 320 and 117 proteins were uniquely identified in early and late ovaries, while 4007 were common in both samples (Fig.7a). Multivariate discriminative analysis indicated a significant separation between the two stages, with PC1 (53.4%) and PC2 (32.5%) contributing to the distinction (Fig.7b). Heat map and hierarchical clustering analysis visually confirmed distinct clusters corresponding to early and late ovaries (Fig.7c). Exploring the biological functions of the identified proteins, we observed a downregulation of apoptosis-related genes and an upregulation of autophagy-related genes in late ovaries compared to early ovaries (Fig.7d). Substantiating the proteomic findings, Western blot analyses validated increased autophagy levels (LC3) in late ovaries compared to early ovaries, suggesting a transition from apoptosis to autophagy during folliculogenesis in goat ovaries, similar to mice (Fig.7e). To further examine the impact of Cathepsin B inhibition, we employed an air-liquid interphase system for the in vitro culture of ovaries. 10 dpp goat ovaries were cultured with or without Cathepsin B and Cathepsin D inhibitors (Fig.7f). After 48 hours, the ovaries were fixed, sectioned, and subjected to immunostaining for the germ cell-specific marker, mouse vasa homologue (MVH), and the guardian of germ cells, TAp63α (Fig.7g). Analysis of follicle counts revealed a 1.5-fold increase in follicles in ovaries treated with Cathepsin B inhibitor compared to the control group, suggesting that Cathepsin B inhibition may protect the ovarian reserve in higher-order vertebrates (Fig.7h).

## Discussion

Folliculogenesis in humans commences with an impressive count of approximately 7 million oocytes, yet only a meager 400 manage to progress to the pre-ovulatory stage over a woman’s entire lifetime. This represents less than 1% of all human oocytes, as the majority succumb to atresia during the folliculogenesis process. These phenomena are universally observed across species. Remarkably, 90-95% of oocytes face elimination during the primordial or primary follicle stage, with the rest undergoing death at the antral follicle stage. The conventional understanding is that oocyte loss occurs in two distinct phases: firstly, postcoital, and secondly, postpartum. Postcoital oocyte loss, attributed to nest breakdown, retrotransposons, and meiotic errors, involves apoptosis and autophagy ^3^. Conversely, meiotically arrested oocytes undergo maturation after birth, progressing through primordial, primary, secondary, preantral, and antral follicles. Notably, the mechanisms behind the demise of primordial and primary follicles still need to be more understood. This study delves into the intricate interplay of various cell death pathways, particularly emphasizing the crucial roles of autophagy and the lysosomal cysteine protease Cathepsin B in preserving the ovarian reserve during the early postpartum stage. This stage marks the second substantial loss of follicles, offering valuable insights into the underlying processes.

### Autophagy and apoptosis equally contribute to maintaining the ovarian reserve

Earlier studies in human ovaries indicate the presence of apoptosis, as observed through TUNEL, with 23.2% and 23.4% positivity in primordial and primary follicles ^31^. Moreover, in mice subjected to chemotherapeutic conditions, inhibiting apoptosis preserves the ovarian reserve, implicating apoptosis in the early loss of ovarian reserve ^32^. However, the involvement of other death pathways remains unclear. Our initial investigation into the regulation of oocyte reserve highlights a delicate equilibrium between apoptosis and autophagy, both crucial processes in ovarian follicle development (Fig.1). Experimental inhibition of apoptosis and autophagy, particularly in early follicles, significantly amplifies the number of ovarian follicles, underscoring their pivotal roles in folliculogenesis (Fig.1 and S1). Simultaneous inhibition proves to be lethal, underscoring the intricate nature of cell death pathways and their role in maintaining the ovarian reserve. Furthermore, our results showcase a significant increase in autophagy levels during folliculogenesis or ageing, establishing a correlation between autophagy and ovarian reserve. Untargeted proteomics analysis unravels the dynamic regulation of apoptosis and autophagy-related genes at different stages of ovarian development, indicating a shift from apoptosis to autophagy during folliculogenesis (Fig.2a-e). This transition is further supported by Western blot analyses, reinforcing the significance of autophagy in preserving the ovarian reserve (Fig.2f-i).

### Cathepsin B in orchestrating oocyte quality and quantity

One of the key findings of the study is the upregulation of lysosomal cysteine proteases, particularly Cathepsin B, which is similar to autophagy during folliculogenesis. Substantial evidence supports the pivotal role of Cathepsin B in programmed cell death. In pathological conditions, there is an increased expression and activity of Cathepsin B, leading to lysosomal membrane permeabilization and subsequent release into the cytoplasm. This release, in turn, triggers diverse programmed cell death pathways, encompassing apoptosis and autophagy ^33^. This study found that inhibiting Cathepsin B leads to a noteworthy increase in oocyte reserves, underscoring its crucial role in preserving ovarian reserve (Fig.3a-d). Genetic requirement studies indicate that Cathepsin B operates within the autophagy pathway to maintain ovarian reserve, signifying its integral involvement in ovarian reserve maintenance via autophagy (Fig.S3b). Moreover, prolonged observations highlight the protective effects of Cathepsin B inhibition, demonstrating sustained preservation of primordial and primary follicles (Fig.S2c). Notably, the study establishes that this protective effect is not detrimental to fertility, as evidenced by the normal reproductive outcomes in treated animals (Fig.S2d-j). The study extends its scope to higher-order vertebrates, showcasing the conservation of the Cathepsin B-dependent mechanism in goat ovaries (Fig.7). This cross-species validation enhances the applicability of our findings, emphasizing the evolutionary significance of this regulatory pathway in maintaining both the quality and quantity of oocytes. While our study utilized chemical inhibitors to assess the impact of Cathepsin B inhibition on ovarian reserve, leaving 1/5th of the protein intact, complete knockout experiments could yield more conclusive results. Previous research has established Cathepsin B’s dispensability for both male and female fertility, making it a promising therapeutic target for ovarian reserve protection ^34^. In addition, recent studies suggest that Cathepsin B inhibitors exhibit therapeutic potential in treating cancers and brain injuries, indicating targeting Cathepsin B may help in several pathological conditions ^35, 36^.

**Fig. 7:**
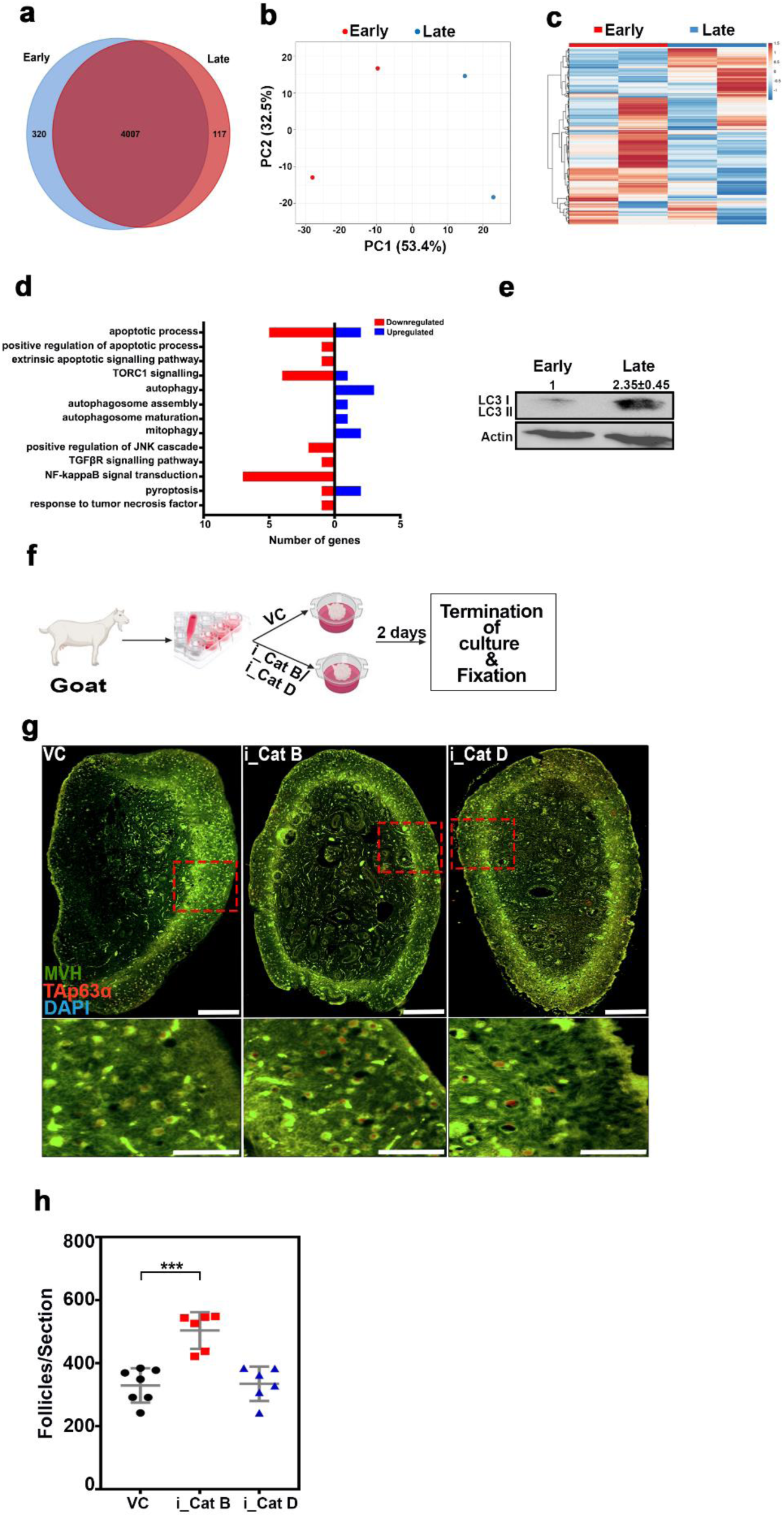
Evolutionary conservation of Cathepsin B dependent oocyte protection. **a,** Venn diagram of early and late goat ovaries proteome. **b**, Principal Component Analysis (PCA) shows the difference between the early and late goat ovaries proteome profiles. **c,** Relative abundance of differential proteins in early and late goat ovaries presented as a heatmap. **d,** Distribution of death pathways related proteins identified in early and late goat ovaries. **e**, Western blot analysis of LC3 and Actin from early and late goat ovaries. **f,** Experimental regime for Ex-vivo goat ovary culture experiments. **g,** VC, i_Cat B and i_Cat D-treated ex-vivo cultured goat ovary sections immunostained for MVH (green), TAp63α (Red), and DNA (blue). **h**, Follicle counts at 48hrs after ex-vivo cultured goat ovary sections. VC, vehicle control; i_Cat B, an inhibitor of Cathepsin B; i_Cat D, an inhibitor of Cathepsin D. The zoomed images below of the ovaries are represented with red squares. \*\*\**P* ≤ 0.0002, *ns ≥* 0.1 unpaired t test. Error bars show mean ± SD. Non-significance P values are not represented in the figure. Scale bars for ovary sections are 200 μm.

### IGF1R links Cathepsin B to mitochondrial dynamics to maintain the quality of the early follicles

This study reveals a novel association between Cathepsin B and IGF1R, highlighting that inhibiting Cathepsin B leads to increased levels of IGF1R (Fig.4e and g). In vitro experiments further illustrate that IGF1R serves as a substrate for Cathepsin B (Fig.4h-m). Exploring the downstream effects, our study elucidates that elevated IGF1R levels impact mitochondrial membrane potential through the transcriptional regulation of genes associated with mitochondrial biogenesis and mitophagy, provides a mechanistic understanding of how Cathepsin B influences oocyte quality (Fig.6c, d, k,l and m). Our findings align with previous research indicating that IGF1 signaling governs mitochondrial dynamics and turnover in cancer cells through the NRF2-BNIP3 pathway ^37, 38^. These results suggest that the protective effects of inhibiting Cathepsin B on ovarian reserve may be attributed to enhanced mitophagy, contributing to the preservation of high-quality oocytes. Then the question is, what role does Cathepsin B play in regulating ovarian reserve? Our findings indicate that elevated oxidative stress in the follicular microenvironment may harm the mitochondria of oocytes (Fig.6i and j). Consequently, these oocytes with damaged mitochondria might experience cell death by suppressing mitophagy through the regulation of IGF1R turnover. A crucial observation is that even though all oocytes may encounter varying levels of oxidative stress, only a select few undergo cell death. This inconsistency could be attributed to differing Cathepsin B and IGF1R levels in the follicular environment (Fig.S5a-f). Our study unveiled variations in the localization of Cathepsin B and IGF1R in different follicles, suggesting that follicles undergoing cell death might exhibit higher Cathepsin B levels. This increase in Cathepsin B might potentially decrease IGF1R levels, subsequently reducing mitophagy and leading to follicular death (Fig.S5a-f). Further research is required to substantiate this hypothesis. In conclusion, this study significantly advances our comprehension of the molecular intricacies involved in maintaining oocyte reserves. The roles identified for autophagy, Cathepsin B, and IGF1R provide valuable insights that may shape the development of innovative fertility treatments and interventions.

## Materials and methods

### 1.1 Animal experiments

The mice were maintained on a 12-hour light-dark cycle with food and water provided ad libitum in accordance with the guidelines set forth by the National Institute of Animal Biotechnology Ethics Committee. C57BL/6 mice were utilized throughout the experiments, except for the evaluation of oocyte and embryo quality, for which FVB mice were employed. In experiments concerning death pathways, 5 postnatal day (dpp) (n = 4 to 7 per inhibitor) were divided into two groups: vehicle control (VC) and inhibitors of apoptosis (i_Apo) [MCE- HY-16658B, (Z-VAD-FMK) 5 mg/Kg body wt.], autophagy (i_Aut) [TCI- M2518, (3-MA) 50 mg/Kg body wt.], pyroptosis (i_Pyr) [Sigma- PHR1690, (Disulfiram) 10 mg/Kg body wt.], and necroptosis (i_Nec) [Sigma- 5303890001, (GSK^’^ 872) 5 mg/Kg body wt.]. The inhibitors were administered intraperitoneally every other day from 5dpp to 10dpp at specified doses. Ovaries were harvested at 10 dpp to maximize the observation of inhibitor effects during the period of higher follicular numbers. For Cathepsin experiments, 10dpp pups (n = 3 to 8 per group) were divided into VC, i_Cat B [SRL- 15358, 37.5 mg/Kg body wt.], and i_Cat D [Sigma- 77170, 20 mg/Kg body wt.] groups. Based on previous studies and lethality assessments, the pups received intraperitoneal injections of Cathepsin B (i_Cat B) or Cathepsin D (i_Cat D) inhibitors. Ovaries were collected on 21dpp, with a subset of i_Cat B pups aged 6 months to evaluate long-term effects. To investigate the interaction between IGF1R and Cathepsin, 10dpp pups (n = 5 to 6 per inhibitor group) were divided into VC, i_IGF1R [Selleck- S7668, 2.5 mg/Kg body wt.], i_Cat B, and i_IGF1R+i_Cat B groups. Intraperitoneal injections of the respective inhibitors were administered every other day from 10dpp to 20dpp, and ovaries were collected at 21dpp. Additional studies involved the administration of i_Cat B to 10dpp pups for JC1, mitotracker, lysotracker, and ROS assessments, with ovaries collected on 21dpp. Endocrine profiling and fertility tests were conducted by injecting i_Cat B into 10dpp pups, while oocyte and embryo assessments were performed on 10dpp FVB mice pups treated with i_Cat B until 20dpp, followed by sacrifice on 45dpp.

### 1.2 Assessment of mitochondrial distribution, membrane potential, ROS, and lysosome activity

The ovaries were delicately trimmed and sliced using a fine 31-gauge needle immersed in M2 media (Sigma- M7167). Subsequently, follicles were carefully isolated under microscopic observation and thoroughly rinsed in media. These follicles were then categorized based on the number of cumulus cell layers they possessed: those with a single layer were denoted early follicles, while those with two or more layers were classified as late follicles. For the assessment of mitochondrial distribution and membrane potential, the follicles were treated with either 2μM of JC-1 (ThermoFisher - M34152) or 40μM of Mito Tracker Green FM (ThermoFisher - M7514) in M2 media at a temperature of 37°C for 30 minutes. Following this incubation period, the follicles underwent thorough washing with M2 media and subsequent rinsing with 1X PBS before being transferred to a dish for immediate microscopic imaging. Excitation at 488nm was utilized for Mito Tracker Green, while JC1 fluorescence was captured at 488nm and 555nm. The fluorescence intensity ratio at 555nm to that at 488nm was computed to determine the membrane potential. To evaluate lysosomal activity, follicles were exposed to 0.2μM lysotracker (ThermoFisher – L7528) for 5 minutes, followed by a series of rinses: three with M2 media and three with 1X PBS. Microscopic images were quickly captured with excitation at 555nm. For the quantification of reactive oxygen species (ROS) levels, follicles were treated with 10μM of 2′,7′ dichlorodihydrofluorescein diacetate (H2DCFDA) (Invitrogen- D399) for 5 minutes, then subjected to similar washing steps before immediate microscopic imaging.

### 1.3 RNA isolation and mRNA expression

Upon sacrificing, ovaries were collected in 1ml of TRIzol (Takara - 9108). Then, they were chopped finely, followed by vortexing and centrifugation at 10000 rpm for 2 minutes at 4^0^C. The supernatant was collected, and RNA was isolated using the phenol-chloroform method. RNA concentration was measured, and cDNA was synthesized according to manufacturer instructions (Takara - 6110a). 100 ng cDNA was used to check the mRNA expression of BNIP3, TFAM, Actin and GAPDH (primers in Table 1) in ovaries by using TB Green syber mix (Takara - RR820A) as per manufacturer instructions. The relative expression of the PCR product was quantified using the (−ΔΔCt) method following Livak’s approach. Statistical significance was established at 0.05% (p<0.05).

**Table 1.**
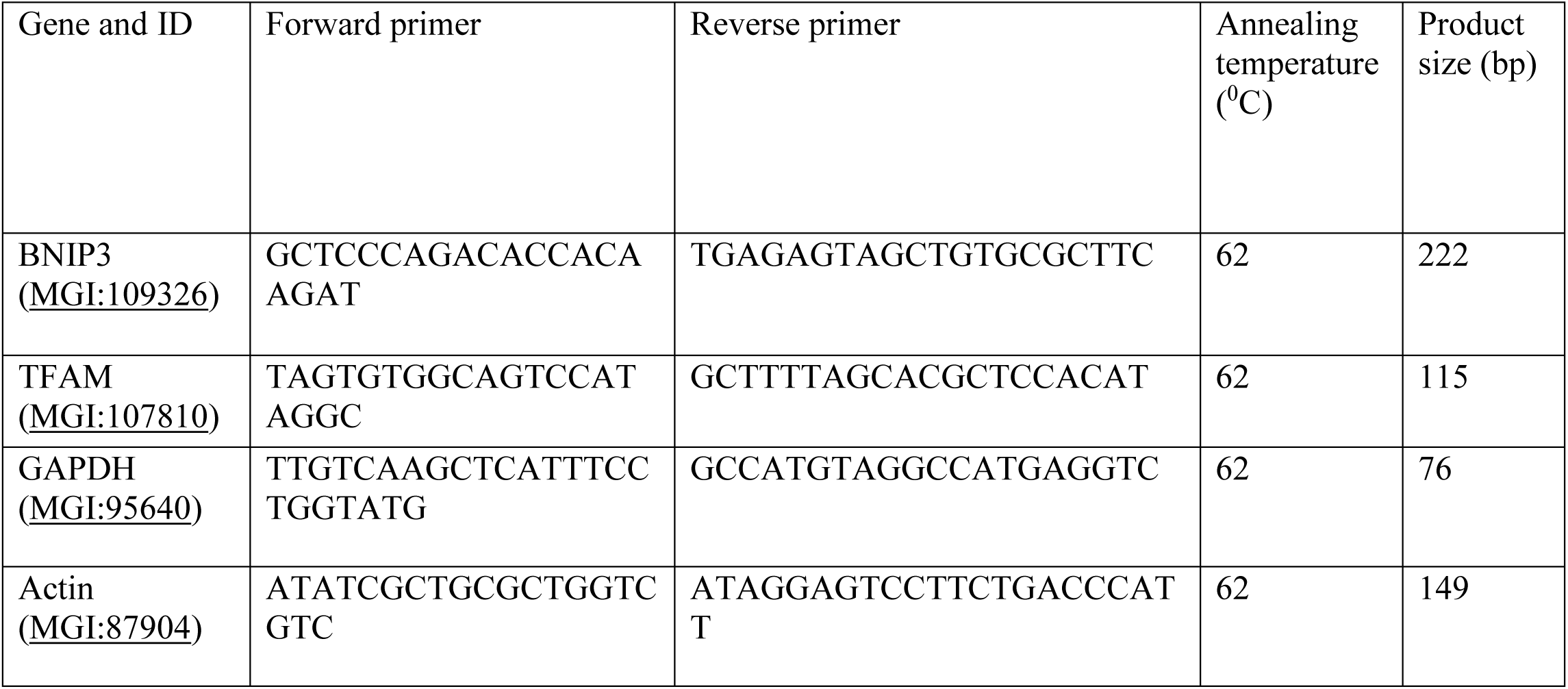

### 1.4 Endocrine Profiling

The serum samples were processed for LH (KLM0573), FSH (K02-0257), AMH (KLM1096) profiling according to manufacturer’s protocol.

### 1.5 Oocyte and Embryo Culture

Ovaries were trimmed and punctured with a 24-gauge needle in M2 media (Sigma- M7167). Under microscopic observation, oocytes were delicately retrieved from the media and thoroughly cleansed through multiple washes. Subsequently, these oocytes were allowed to mature within the same M2 media within the controlled environment of a CO2 incubator set to 90-95% humidity and 37°C. After 16 hours of incubation, the oocytes were examined for signs of maturation (polar body extrusion and DNA staining). To collect embryos, animals underwent super-ovulation through injections of PMSG and HCG. Once primed, the ovaries and the oviducts from plugged females were carefully dissected. The oviducts underwent trimming, and swollen ampullas were gently punctured using a 24-gauge needle within M2 Media. Embryos, still encased in cumulus complexes, were washed twice with M2 media and once with Embryomax HTF media (Sigma- MR-070-D). Following this, the embryos were cultured using the same Embryomax HTF media within a 4-well IVF dish (Thermo-144444), all within the controlled conditions of a CO2 incubator set to 90-95% humidity and 37°C. After 4 days, the embryos were microscopically observed for signs of maturation.

### 1.6 Fertility Studies

Individual adult female mice, treated and untreated, were housed separately with untreated males to evaluate fertility. If a female failed to conceive within a month, the male partner was replaced. The females underwent plug-checking for a week or until successful mating with a new male, confirmed by the presence of a mating plug.

### 1.7 Goat Ovary Culture

Ovary samples from the Osmanabadi goat breed obtained one day after birth, were collected from the slaughterhouse, and immediately immersed in a warm saline solution (32-37°C) fortified with penicillin–streptomycin antibiotics. Within an hour of collection, the samples were transported to the laboratory for further processing using an embryo carrier. Upon arrival, the ovaries were carefully trimmed and placed on a filter strip (782810). This filter was then floated on a nutrient medium consisting of TCM199 (Sigma - M2154) supplemented with 10% goat follicular fluid, 1× penicillin–streptomycin (Sigma -P4333), and 3% fetal calf serum. The culture plates (782891) containing the samples were then placed in a controlled environment containing a CO2 incubator set at 37°C with 90-95% humidity. After 2 hours of culture, either 50μM of i_Cat B or 1μM of i_Cat D, along with their respective controls, were introduced into the culture. The culture was allowed to continue for a total of 48 hours. Upon completion of the culture period, the ovaries were fixed in 10% formalin and underwent further processing for sectioning. All experiments were conducted with three independent biological replicates to ensure the reliability and consistency of the results.

### 2.1 In vitro Cathepsin B reaction

For in vitro reaction,1μg of purified IGF1R (R&D systems- 305-GR) was incubated with assay buffer 30μl [25 mM MES (pH 5)] in the presence or absence of purified Cat B [R&D systems- 953-CY, (500 ng)]. After 2 hrs incubation at 37°C, the reaction was kept at -20 °C to quench the reaction.

### 3.1 MS sample preparation

For whole-cell untargeted proteomics of ovarian samples, initially, ovaries were lysed using a solution containing 1% Sodium Deoxycholate (SDC) prepared in 50 mM Ammonium Bicarbonate Buffer (AMBIC). Following this, the protein concentration in the lysate was determined using the Bradford assay, and equal amounts of protein from each sample were selected for subsequent processing. Samples subjected to in vitro reactions were directly processed after the reaction. The process began with treating the samples with 200 mM Dithiothreitol (DTT) to reduce disulfide bonds, which was carried out at 57°C for 1 hour. Subsequently, alkylation was performed using 200 mM Iodoacetamide (IAA) at room temperature in the dark for 1 hour. The samples were then digested using a combination of trypsin and LysC enzymes at 37°C for 16 hours. To remove Sodium Deoxycholate (SDC) from the trypsin-digested samples, 20% formic acid was added to precipitate it. The resulting mixture was then passed through a 10 kDa filter to eliminate undigested proteins. Further purification was achieved by desalting the samples using a C18 spin column following the manufacturer’s instructions. The purified samples were subsequently vacuum-dried. Further, the dried samples were reconstituted in 0.3% formic acid. Finally, 1 μg of the peptide sample was injected into the mass spectrometer for analysis. All experiments were done with two to three independent biological replicates.

### 3.2 Mass Spectroscopy Data Acquisition

The proteome was analysed by using the UltiMate 3000 RSLCnano system coupled with the high-resolution Q Exactive HF mass spectrometer (Thermo Fisher Scientific). In brief, the full MS scans were carried out by using a resolution value of 60000, AGC target value of 1 × 10^6^ and acquisition range of 375–1600 *m/z* with a maximum injection time of 60 ms. The top 25 precursors were selected for the fragmentation. The MS2 acquisition was performed through a resolution of 15,000, AGC target value of 1 × 10^5^, a maximum injection time of 100 ms, an isolation window of 1.3 m/z, and a fixed first mass at 100 *m/z*. A nonlinear gradient (flow rate of 0.300 μL/min for 180 min) of solvents using 5% of 80% acetonitrile/0.1% formic acid as solvent B and 95% of 0.1% formic acid as solvent A was preferred for eluting the peptides.

### 3.3 Mass spectroscopy data analysis

The peptides and proteins were identified by using the software Proteome discoverer v2.5 (Thermo Fisher Scientific, San José, CA, USA) with UniProtKB/Swiss-Prot: P08069.1, UniProtKB Mice (UP000000589) and Goat (UP000291000) database.

### 3.4 Parameters for peptide and protein search

We selected the dynamic and static modifications of oxidized methionine residues and carbamidomethylation of cysteine residues as our prime search parameters. In addition, we considered the following values of 2,144 and 6 for most missed cleavage, maximum and minimum peptide length respectively. Proteins and peptides with < 1% and < 5% FDA confidence were filtered. The identification of proteins and peptides were based on the 10 ppm and 0.02 Da fragment and precursor mass tolerances. The ‘Minora Feature Detector’, ‘Precursor Ions Quantifier’ and ‘Feature Mapper’ nodes workflow were used for the Label-free quantification (based on peptide signals) to identify proteins or peptides ^39^.

### 3.5 Bioinformatics and Statistical Analysis

Following the label-free quantification, protein intensities (abundance values) were normalized by log2 transformation. The inter-variation and intra-variation between different samples were determined by using the Venn diagram and PCA respectively. Heat map and PCA graphs were generated by using Clutvis (http://biit.cs.ut.ee/clustvis/.). Gene ontology (GO) comparative analysis was performed with Funrich v3.1.3 and ShinyGO v0.741 software ^40, 41^.

### 4.1 Cell culture

Human lung adenocarcinoma cell line H1299 purchased from ATCC (ATCC - CRL-5803) was grown in RPMI1640 (Gibco - 11875085), 10% FBS (Gibco - 10270106) and 1× penicillin– streptomycin (Sigma -P4333) at 37 °C and 5% CO2. For Cat B overexpression, the cells were transfected with hCathepsin B (add gene-11249) plasmid using Lipofectamine™ 3000 transfection Reagent (Thermo- L3000008) following manufacturer’s protocol. The cells were harvested after 24hrs of transfection. To inhibit Cathepsin B, cells were cultured with or without i_Cat B (50μM) for 4hrs and then harvested. Harvested cells were then centrifuged at 15000rpm at 4^0^C for 10 minutes to collect the pellets. The pellets were resuspended in lysis buffer (25mM Tris-HCL, pH-8.0; 250mM NaCl; 1% Triton-X-100; 1% SDS; 2mM MgCl2; PMSF; Protease inhibitor) followed by sonication and centrifugation at 12,000 rpm at 4^0^C for 10mins. Supernatants were then boiled at 100^0^C for 5mins with 1X Laemmle buffer and loaded in SDS-PAGE to check the protein of interest.

### 5.1 Immunohistochemistry and Immunofluorescence

The ovaries were first fixed in 10% formalin for 2-4 days. Fixation was followed by a gradient treatment of 10%, 20%, and 30% sucrose up to 3 days depending on the size of the ovary. Then 5 µm thin cryosections were obtained by the help of Cryostar NX50 and placed on glass slides with positive surface coating. The slides were then deparaffinized for further staining processes. Next the slides were treated at 95°C for 60 minutes in antigen retrieval buffer comprising 10 mM Sodium Citrate and 0.05% Tween 20 for antigen retrieval necessary in further immunofluorescence staining. After cooling to room temperature under running water, the tissue sections on slides underwent two times dH2O based washing for 5 minutes each followed by rinsing with 1XTBST-0.1% Tween 20 for 1 minute. The tissue sections were then blocked with M.O.M (BMK-2202) for 1 hour at room temperature, followed by two rounds of blocking with ADB for 15 minutes each. The sections were incubated overnight in primary antibodies (Rabbit anti-TAp63 (BS1279, 1:500), Mouse anti-MVH (ab27591, 1:500)). Following incubation, slide containing sections were washed two times with 1XTBST-0.1% Tween 20 in 10 minutes interval. Then slides were blocked with ADB twice for 15 minutes each. Afterwards, the slides were incubated with corresponding secondary antibodies (Goat anti-rabbit 555 (A32732, 1:2000), Goat anti-mouse 488 (A32723, 1:2000)) at 37°C for 1 hour in the dark. Later, the slides were washed thrice with 1XTBST-0.1% Tween 20 for 5 minutes each and stained with 0.01% DAPI for 2 minutes. The slides were then air dried after a minute of dH2O washing. For DAB staining, after blocking with ADB, slides were incubated with secondary antibody Goat anti-rabbit biotin (B-2770, 1:1000) at 37°C for 1 hour in the dark. This was followed by three times washing with 1XTBST-0.1% Tween 20 at 5 minutes interval. Subsequently, slides were processed according to the manufacturers protocol i.e., the VECTASTAIN ABC-HRP kit (PK-6100) and DAB substrate kit (SK-4100). The slides were the dried and mounted with DPX, and images were captured under the microscope.

### 5.2 Imaging and quantification

Immuno-stained slide images were captured by the help of Zeiss Axio scope VII microscope with 10×, 20× Plan Apochromat 0.45 NA, or 63×Plan Apochromat 1.4 NA objectives and EXFO X-Cite metal halide light source with a Hamamatsu ORCA-ER CCD camera. The acquired images were processed by the Zen software. In the ovarian samples, we examined every fifth section for different types of follicles: primordial, primary, secondary, preantral and antral. A single layer of squamous or cuboidal epithelial cell containing follicles were considered as primordial or primary follicles, respectively. In a similar fashion, follicles surrounded by two or more layers of cells were considered as secondary follicles. Follicles with early antrum formation were counted as preantral follicles, and completely formed antrum containing follicles were considered as antral follicles. All quantitative analyses were executed by two observers where the second observer was blindfolded in the examined group.

### 6.1 Western blotting

The ovaries were minced and lysed in a lysis buffer containing 25 mM Tris-HCL (pH 7.5), 150 mM NaCl, and 1% Triton-X-100, supplemented with PMSF and Protease inhibitor. Next, the mixture was homogenized, sonicated, and centrifuged at 12,000 rpm for 10 minutes at 4°C. The supernatant was collected carefully and stored at -80°C for further analysis. The protein concentration was determined by using Bradford assay kit (Takara). For the vehicle control and inhibitor-treated or 5, 10 20 dpp ovarian samples equal amounts of protein were loaded. The proteins were separated by SDS-PAGE and then transferred onto a nitrocellulose membrane. The membrane was then blocked with 5% skimmed milk for 1 hour at room temperature, followed by overnight incubation at 4°C with primary antibodies. The primary antibodies used were as follows: Mouse anti-γH2A.X (05-63-I, 1:1000), Rabbit anti-Cleaved Caspase 3 (D175, 1:1000), Rabbit anti-LC3 (NB100-2220, 1:1000), Rabbit anti-P62 (5114T, 1:1000), Rabbit anti-TAp63α (BS1279, 1:1000), Rabbit anti-MVH (ab13840, 1:1000), Mouse anti-Cathepsin B (C6243, 1:500), Rabbit anti-Cathepsin D (E7Z4L, 1:1000), Mouse anti-ATG7 (MAB6608, 1:1000), Mouse anti-IGF1 (sm 1.2, 1:1000), Rabbit anti-IGF1R (9750, 1:1000), Rabbit anti-phospho-IGF1R (3021, 1:1000), Rabbit anti-phospho-AKT (4060, 1:1000), Rabbit anti-phospho-mTOR (5536, 1:1000), Mouse anti-Tubulin (SC 53646, 1:1000), Mouse anti-β- Actin (SC-47778, 1:1000). After the overnight incubation, the blots were washed thrice at an interval of five minutes with 1X TBST consisting 0.3% Tween 20 and then incubated with corresponding secondary antibodies labelled with HRP (Goat anti-Mouse IgG HRP,31430; Goat anti-Rabbit IgG, 31460) for 2 hours at room temperature. After three rounds of washing with 1X TBST with 0.3% Tween 20 and a final wash with 1X TBS, the blots were developed using chemiluminescence. The Western blot signals were quantified using Image J software.

### Methods contact

Further information and requests for resources and reagents should be directed to the lead contact H.B.D. Prasada Rao

## ACKNOWLEDGMENTS

We thank the NIAB core Microscope and small animal facility. A.M. was supported by CSIR SRF. A.K. was supported by DBT SRF. L.K.S was supported by DBT SRF. R.B. was supported by UGC SRF. A.K., P.K., was supported by UGC JRF. This work was supported by ICMR grant (2020-4310), DBT grant (BT/PR46563/AAQ/1/868/2022) awarded to H.B.D.P.R.

## AUTHOR CONTRIBUTIONS

A.M., and H.B.D.P.R. conceived the study and designed the experiments. A.M., A.K., A.K., P.K., L.K.S, R.B., and H.B.D.P.R. performed the experiments and analyzed the data. H.B.D.P.R., and A.M. wrote the manuscript with inputs and edits from all authors.

## Conflict of interest

The authors declare no competing interests. None of the material reported in this manuscript has been published or made available online, nor is it under consideration elsewhere.

## Data availability

The data that support the findings of this study are available from the corresponding author upon request.

## Supplementary Figure Legends

**Fig. S1:**
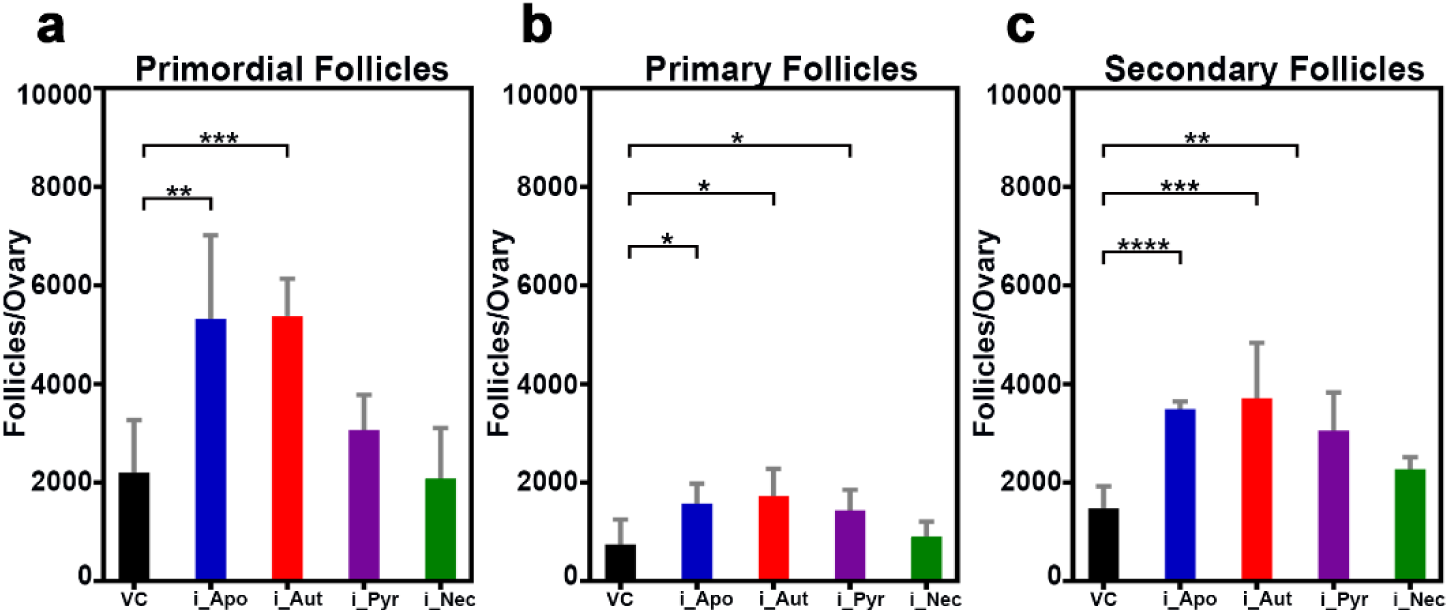
**a, b and c,** Quantification of primordial, primary, and secondary follicles at 10 dpp from VC, i_Apo, i_Aut, i_Pyr, and i_Nec -treated mice ovaries. VC, vehicle control; i_Apo, an inhibitor of apoptosis; i_Aut, an inhibitor of autophagy; i_Pyr, an inhibitor of pyroptosis; i_Nec, an inhibitor of necrosis. \*\*\*\**P* ≤ 0.0001, \*\*\**P* ≤ 0.001, \*\**P* ≤ 0.004, \**P* ≤ 0.04, *ns ≥* 0.1 unpaired t test. Error bars show mean ± SD. Non-significance P values are not represented in the figure.

**Fig. S2:**
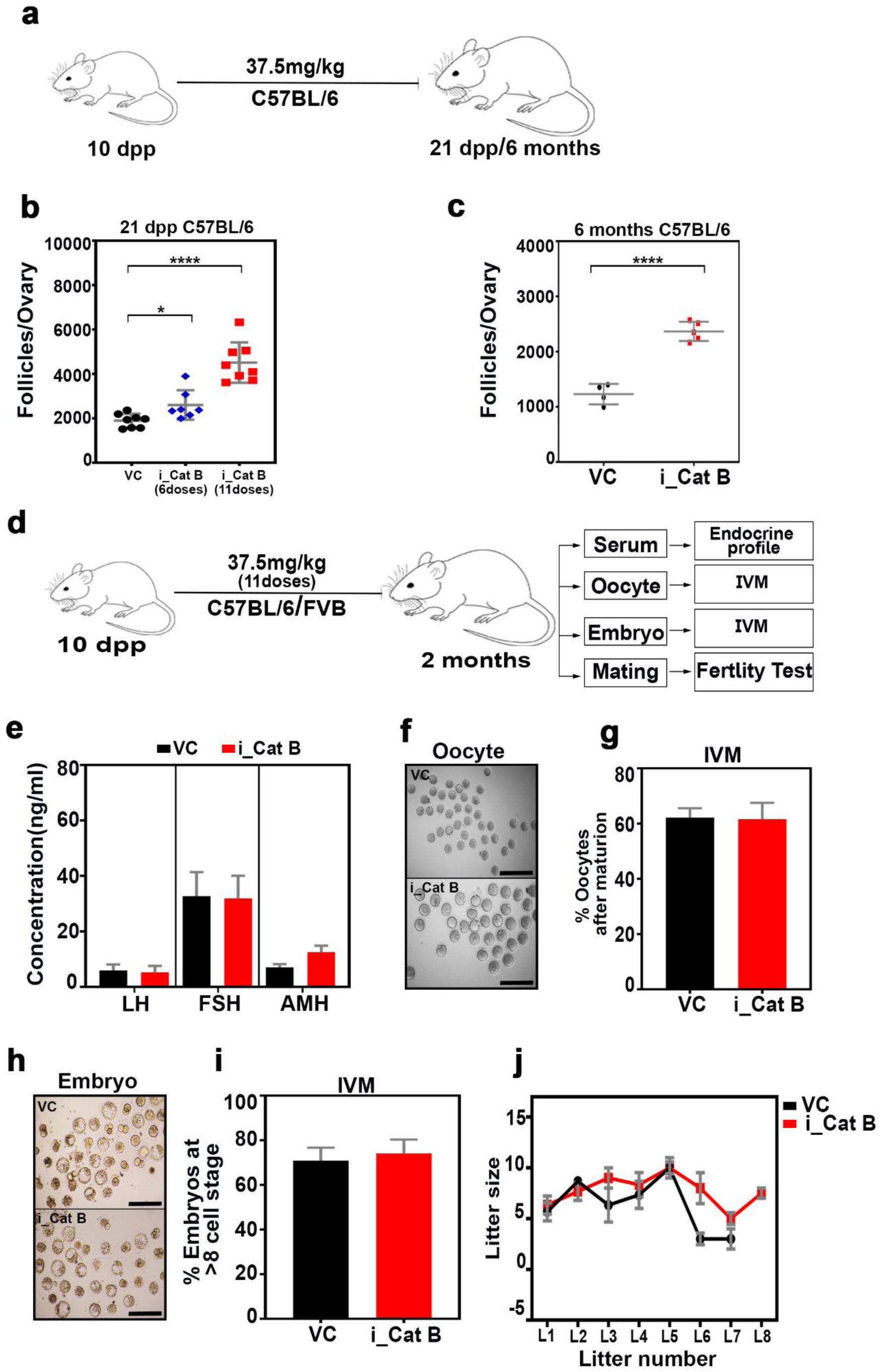
**a,** Experimental regime for i_Cat B intraperitoneal injection with different doses and in aged mice ovaries. **b,** Follicle counts at 21 dpp with different doses. **c**, Follicle counts at 6 months of 11-dose treatment of VC and i_Cat B mice ovaries, respectively. **d**, Experimental regime for i_Cat B intraperitoneal injection followed by endocrine profiling, oocyte, embryo quality assessment and fertility test experiments. **e**, Quantification of LH, FSH and AMH in the serum from VC and i_Cat B -treated females. **f**, Bright field oocyte maturation images. **g**, Quantification of in-vitro oocyte maturation from VC and i_Cat B -treated females. **h**, Bright field embryo development images. **i**, Quantification of in-vitro embryo development from VA and i_Cat B -treated females. **j**, Fertility performance of VC and i_Cat B -treated females. VC, vehicle control; i_Cat B, an inhibitor of Cathepsin B. \*\*\*\**P* ≤ 0.0001, \*\**P* ≤ 0.001, \**P* ≤ 0.02 *ns ≥* 0.5 unpaired t test. Error bars show mean ± SD. Non-significance P values are not represented in the figure. Scale bars are 20 μm.

**Fig. S3:**
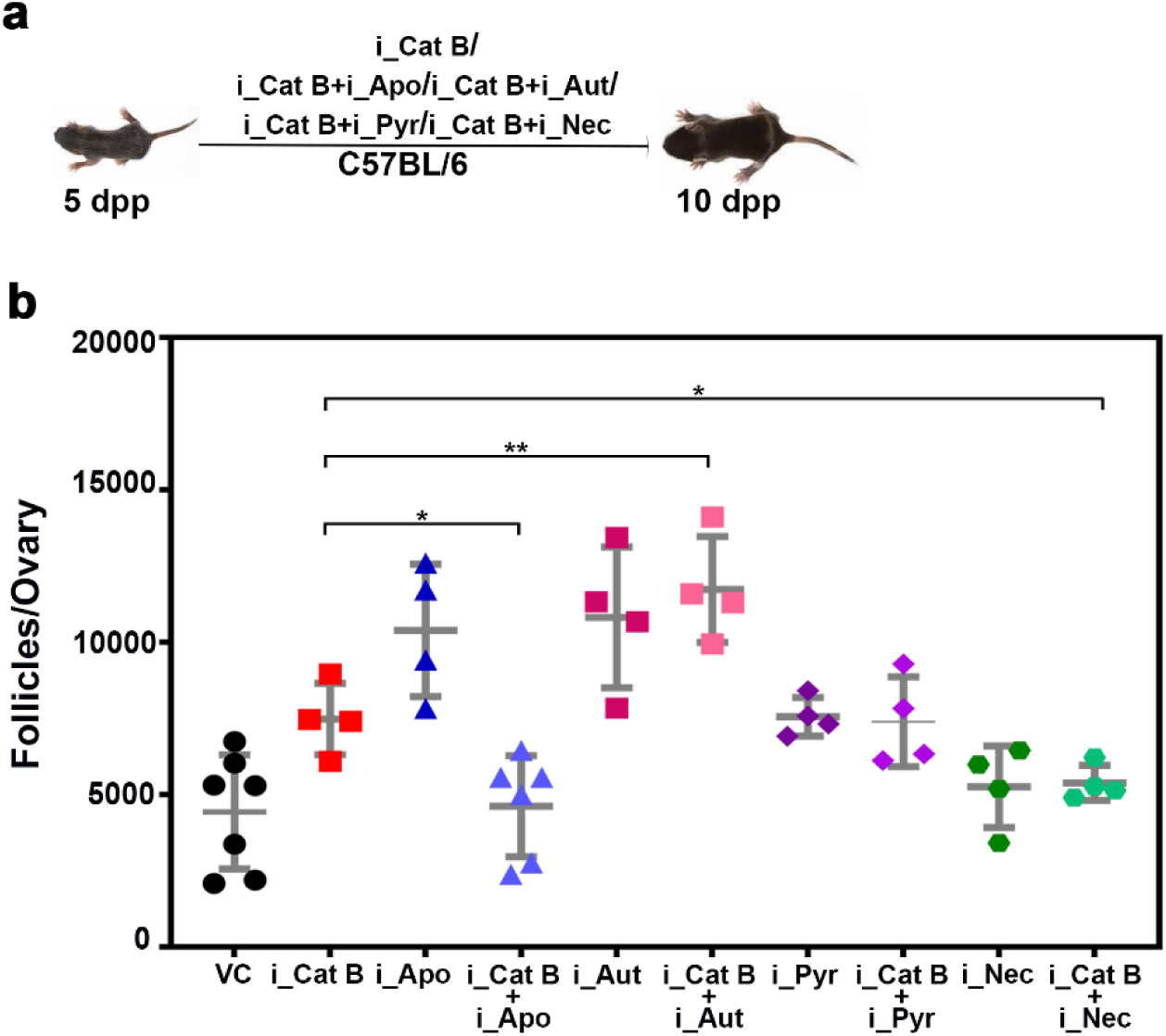
**a,** Experimental regime for i_Cat B intraperitoneal injection with different death pathways inhibitors. **b,** Follicle counts at 10 dpp from VC and i_Cat B alone or along with i_Apo, i_Aut, i_Pyr, and i_Nec - treated females. VC, vehicle control; i_Cat B, an inhibitor of Cathepsin B; i_Apo, an inhibitor of apoptosis; i_Aut, an inhibitor of autophagy; i_Pyr, an inhibitor of pyroptosis; i_Nec, an inhibitor of necrosis. \*\**P* ≤ 0.006, \**P* ≤ 0.01, *ns ≥* 0.5 unpaired t test. Error bars show mean ± SD. Non-significance P values are not represented in the figure.

**Fig. S4:**
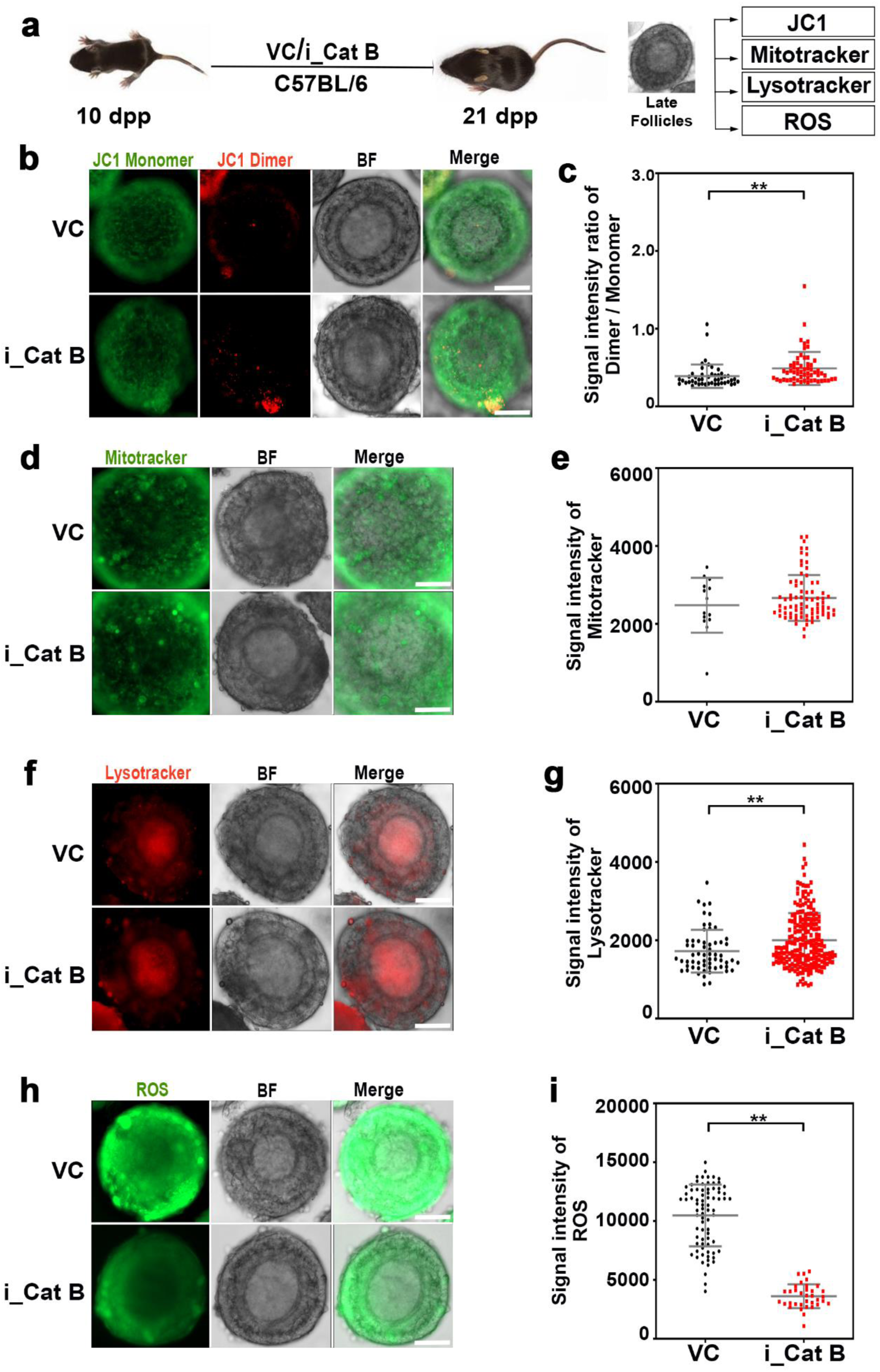
**a,** Experimental regime for intraperitoneal injection of VC and i_Cat B -treated ovaries followed by JC1, mitotracker green, lysotracker red, and ROS experiments. **b,** Late follicles from VC and i_Cat B -treated ovaries were stained for JC1 and imaged at 488nm for JC-1 monomers and 565nm for dimers. **c,** Quantification of the signal intensity ratio of Dimer/Monomer. **d,** Late follicles from VC and i_Cat B -treated ovaries were stained for mitotracker. **e,** Quantification of the signal intensity of mitotracker. **f,** Late follicles from VC and i_Cat B -treated ovaries were stained for lysotracker. **g,** Quantification of the signal intensity of lysotracker. **h,** Late follicles from VC and i_Cat B -treated ovaries were stained for ROS. **i,** Quantification of the signal intensity of ROS. VC, vehicle control; i_Cat B, an inhibitor of Cathepsin B. \*\*\*\**P* ≤ 0.0001, \*\**P* ≤ 0.07, *ns ≥* 0.1 unpaired t test. Error bars show mean ± SD. Non-significance P values are not represented in the figure. Scale bars for follicles are 10 μm.

**Fig. S5:**
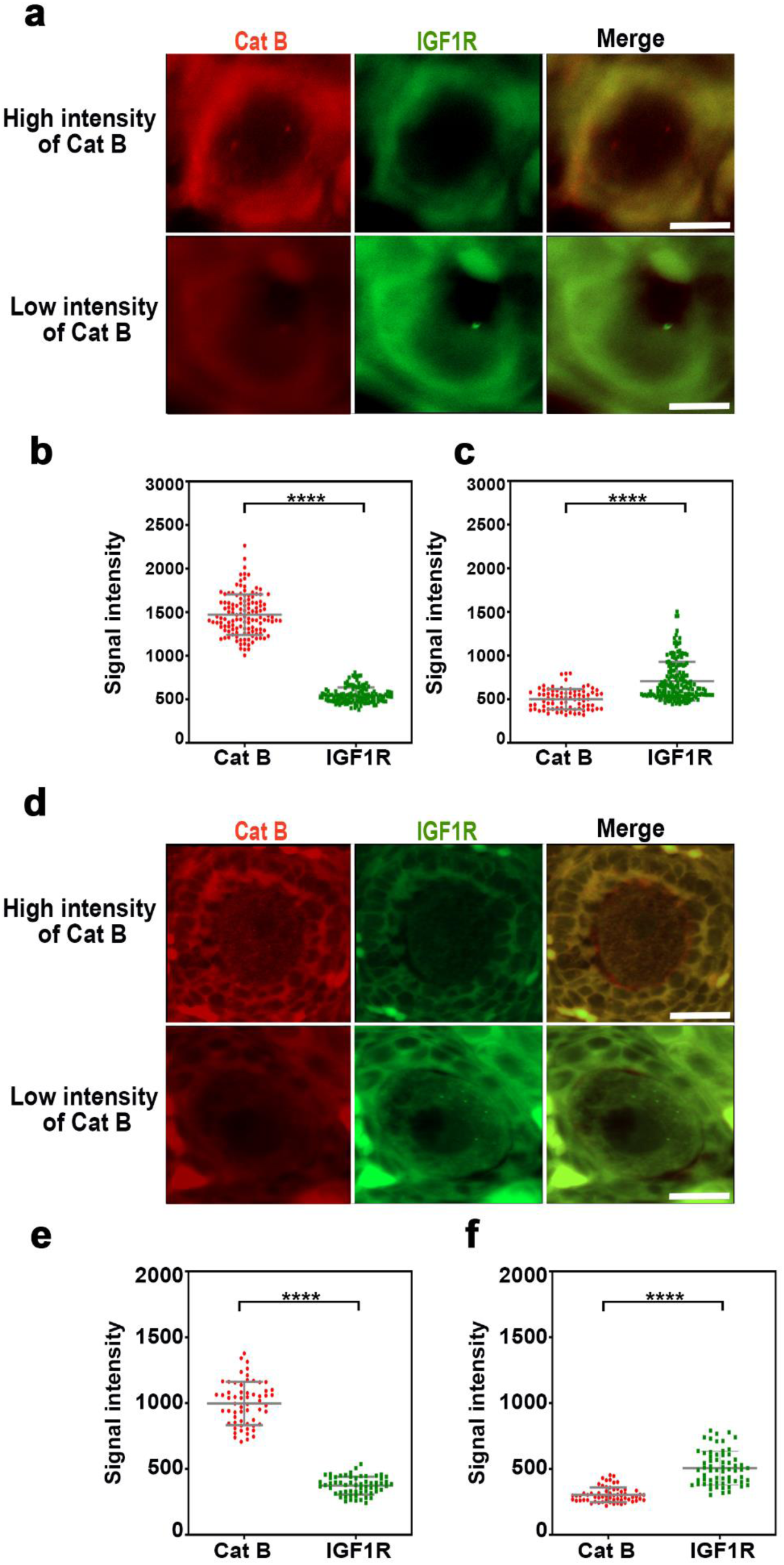
**a,** Representative images of early follicles from 10 dpp mice ovary immunostained for Cathepsin B (Red), IGF1R (green) and DNA (blue). **b and c,** Quantification of the signal intensity of Cathepsin B and IGF1R of early follicles at high and low intensity of Cathepsin B respectively. **d,** Representative images of late follicles from 10 dpp mice ovary immunostained for Cathepsin B (Red), IGF1R (green) and DNA (blue). **e and f,** Quantification of the signal intensity of Cathepsin B and IGF1R of late follicles at high and low intensity of Cathepsin B respectively. Cat B, Cathepsin B. \*\*\*\**P* ≤ 0.0001, unpaired t test. Error bars show mean ± SD. Scale bars for follicles are 20 μm.

**Fig. S6:**
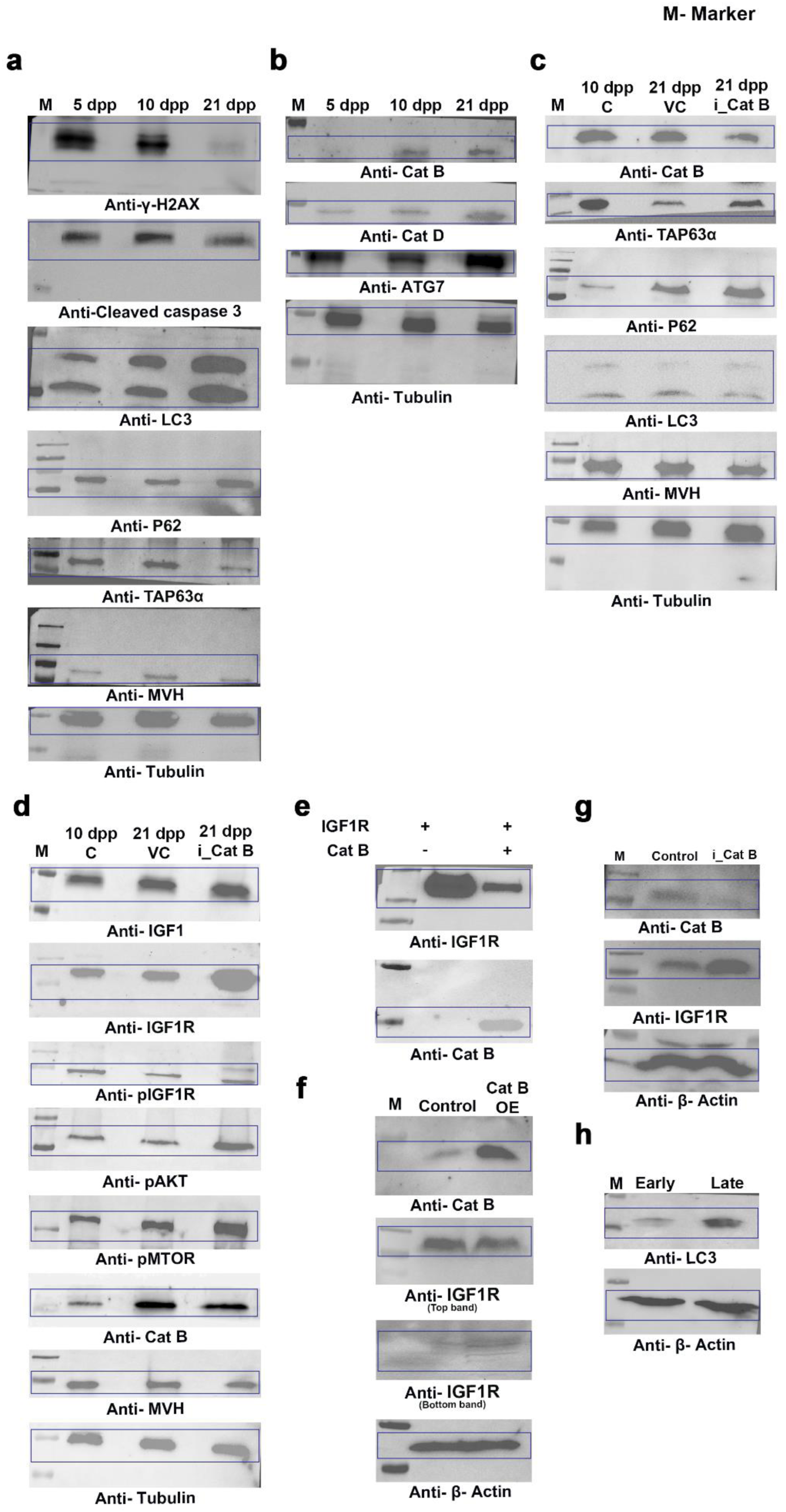
**a-h,** Whole western blots.

## Notes

### Competing Interest Statement

The authors have declared no competing interest.

